# Uncovering the hidden threat: single-organoid analysis reveals clinically relevant treatment-resistant and invasive subclones in pancreatic cancer

**DOI:** 10.1101/2023.02.27.530080

**Authors:** Maxim Le Compte, Edgar Cardenas De La Hoz, Sofía Peeters, Felicia Rodrigues Fortes, Christophe Hermans, Andreas Domen, Evelien Smits, Filip Lardon, Timon Vandamme, Abraham Lin, Steve Vanlanduit, Geert Roeyen, Steven Van Laere, Hans Prenen, Marc Peeters, Christophe Deben

## Abstract

Pancreatic ductal adenocarcinoma (PDAC) is one of the most lethal diseases, characterized by a treatment-resistant and invasive nature. In-line with these inherent aggressive characteristics, only a subset of patients show a clinical response to the standard of care therapies, thereby highlighting the need for a more personalized treatment approach. In this study, we comprehensively unraveled the intra-patient response heterogeneity and intrinsic aggressive nature of PDAC on bulk and single-organoid resolution. We leveraged a fully characterized PDAC organoid panel (N=8) and matched our artificial intelligence-driven, live-cell organoid image analysis with retrospective clinical patient response. In-line with the clinical outcomes, we identified patient-specific sensitivities to the standard of care therapies (gemcitabine-paclitaxel and FOLFIRINOX) using a growth rate-based and normalized drug response metric. Moreover, the single-organoid analysis was able to detect resistant as well as invasive PDAC organoid clones, which was orchestrates on a patient, therapy, drug, concentration and time-specific level. Furthermore, our *in vitro* organoid analysis indicated a strong correlation with the matched patient progression-free survival (PFS) compared to the current, conventional drug response readouts. This work not only provides valuable insights on the response complexity in PDAC, but it also highlights the potential applications (extendable to other tumor types) and clinical translatability of our approach in drug discovery and the emerging era of personalized medicine.

## 1. Introduction

Pancreatic ductal adenocarcinoma (PDAC) is a rapidly progressing and usually fatal disease with a 5-year overall survival rate of less than 10% ^1,2^. Despite significant efforts to improve the clinical outcome, current standard of care treatments show only limited efficacy in both locally advanced and metastatic PDAC patients^3-5^. Numerous studies have shown that this poor outlook can be attributed to the extensive intratumoral clonal heterogeneity, patient-specific transcriptional plasticity, and the intrinsic invasive behavior of PDAC tumor cells^6-8^. Given the patient-specific signatures and aggressiveness of this malignancy, it is unquestionably evident that innovative and more personalized approaches are paramount to treat PDAC patients and improve their quality of life.

Over the past years, patient-derived tumor organoids emerged as promising cancer models for personalized medicine, since they preserve the clonal heterogeneity, mutational landscape, and histological architecture of the originating tumor tissue^9-11^. Furthermore, recent studies provided first evidence that tumor organoids responded similarly to the corresponding patient when treated with the same standard of care therapy. However, these retrospective clinical studies were only able to predict clinical responses in a subset of patients^12,13^. One major factor driving this limitation is that current readouts only extract a fraction of the information that tumor organoids could potentially provide. The current gold-standard analysis method for organoid-based research is the ATP-based endpoint assay, CellTiter-Glo 3D^14,15^. Although this viability assay has been extensively used in numerous studies, it fails to account for the heterogeneity of tumor organoids by relying on a bulk lysis approach^16^. Moreover, it has been shown that intracellular ATP levels dynamically change upon treatment with chemotherapeutics (e.g. senescent vs non-senescent cells), which could further bias this readout^17-20^. Taken together, the current methods compress the highly-dimensional tumor organoid model into an oversimplified, single-layered readout, which can result in misleading clinical interpretation and translatability. Therefore, we hypothesize that by incorporating higher-dimensional analysis methods on a single-organoid resolution, we will further unlock the predictive performance of tumor organoids as ‘patient-in-the-lab’ models for guiding drug development and clinical decision in personalized medicine.

In this study, we combined a normalized drug screening metric (NDR) with the dynamic quantification of single organoid responses to evaluate drug responses (standard of care) in patient-derived PDAC organoids, using our software platform (Orbits)^21,22^. This enabled us to study patient-specific and intratumoral subclonal sensitivities to different standard of care therapies *ex vivo*, which was validated with matched clinical patient response to therapy – progression free survival (PFS). Our data not only highlight the inter-patient heterogeneity in terms of therapy response, but also shed light on the presence of treatment-resistant and invasive subclones that could drive disease progression. Consequently, our approach also enabled us to increase the clinical translatability of organoid analysis by more accurately recapitulating and measuring the complexity of human tumors in matched PDAC organoids. This method will inevitably be of tremendous value for both the development of novel therapeutics and the guiding of more informed clinical decision in the rising era of personalized medicine.

## 2. Results

### Selected panel of PDAC organoids shows a distinct phenotypic, molecular and clinical landscape

To capture the inter- and intra-patient heterogeneity, we selected a panel of 8 representative human PDAC organoid lines based on the following distinct features: morphology, molecular landscape, and clinical characteristics (*in vivo* response, treatment, progression-free survival). All 8 PDAC organoid lines were derived from primary tumor samples taken during surgery. Hematoxylin and eosin (H&E) staining and brightfield image analysis showed high phenotypic variations between the PDAC organoids (baseline and treated) (Fig. 1a, Extended Data Fig. 1a). In-line with previous studies, we identified both cystic/hollow-like (thin layered epithelium) and solid-like (lobular with pleiomorphic cells) morphologies within our patient cohort. Moreover, as growth rates have been reported to potentially vary between individual patients^23^, the baseline PDAC organoid growth rates were also monitored for 5 days, using live-cell imaging. Interestingly, a high degree of growth rate variation between the 8 PDAC organoid lines was revealed, and upon further observation, solid-like PDAC organoids (PDAC068, PDAC070, PDAC082 and PDAC087) appeared to exhibit slower growth patterns compared to the cystic-like organoids (PDAC002, PDAC044, PDAC052 and PDAC060) (Fig. 1a).

**Fig. 1.**
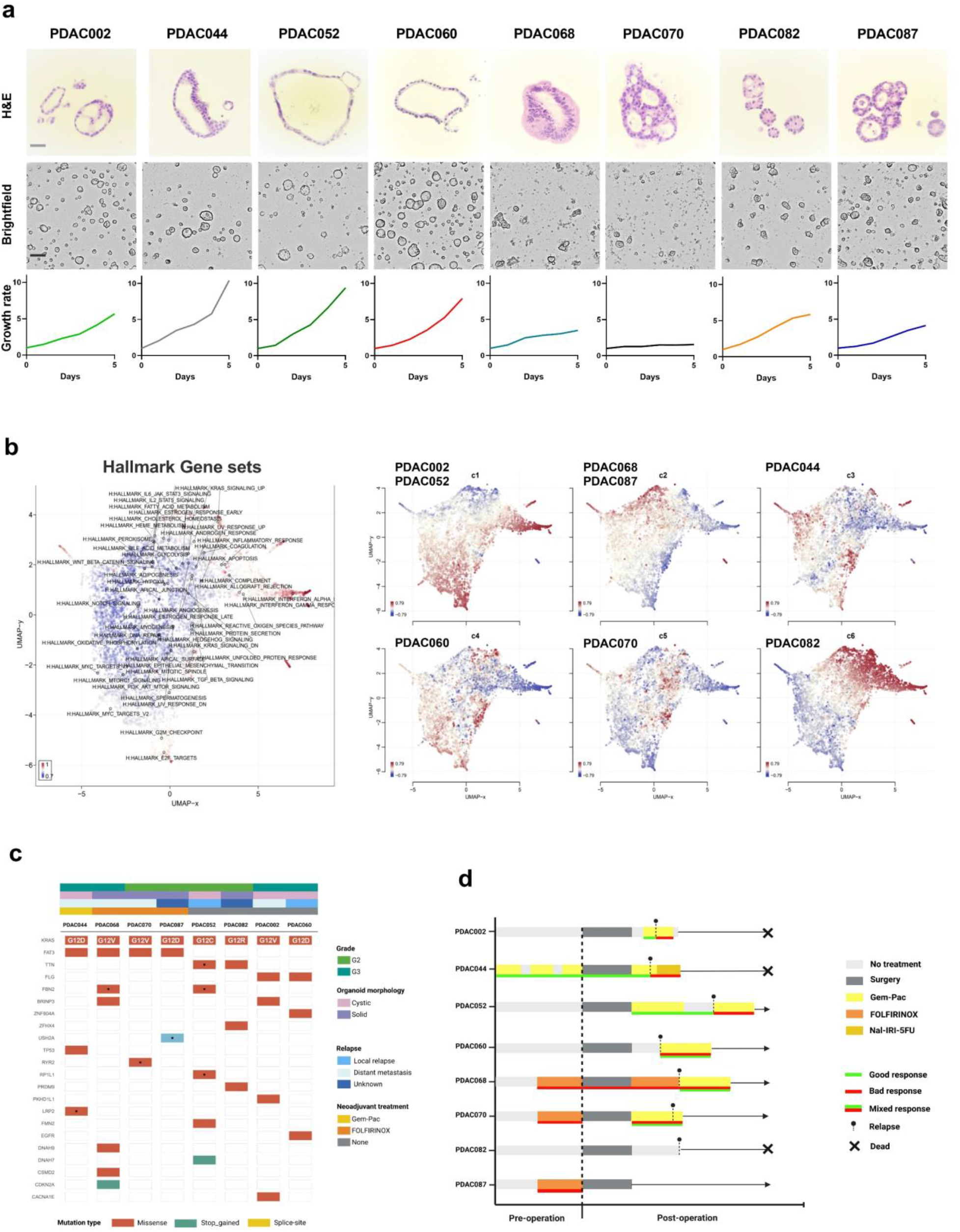
PDAC organoid cohort shows distinct morphological, clinical and molecular features. **a**. Representation of the H&E and brightfield images combined with the associated growth rates of the included PDAC organoids (grey scale bar=30 μm and black scale bar=100 μm). **b**. Uniform Manifold Approximation and Projection (UMAP) visualization of the top 50 differently expressed Hallmark gene sets highlighting the patient-specific transcriptional profiles. **c**. Oncoprint illustrating the key somatic alterations with the annotated clinical information (Grade, Morphology, Relapse and Neoadjuvant treatment). **d**. Timeline of the clinical course showing the treatment, response and relapse/dead of each individual patient. Gem-Pac: gemcitabine/nab-paclitaxel. Good response: tumor regression or stable disease, bad response: no tumor regression and disease progression and mixed response: minimal regression/disease control of metastases but no response at the primary tumor.

The observed morphological and growth rate differences likely stem from distinct molecular landscapes, and therefore, we performed RNA-seq analysis to delineate the different signatures. The differentially expressed gene and Uniform Manifold Approximation and Projection (UMAP) analysis revealed 6 individual clusters (c1-c6) with distinct transcriptional characteristics, which further highlights the inter-patient heterogeneity in PDAC (Fig. 1b, Extended Data Fig. 1b-c). It is also of interest that, compared to the fast cycling PDAC organoids, the solid-like PDAC organoids (PDAC068, PDAC070, PDAC082, and PDAC087) have a reduction in three hallmark cell proliferation-related gene sets (G2M Checkpoint, E2F Targets and mitotic spindle), which can be linked back to their intrinsic slower growth rates (Fig. 1a,b, Extended Data Fig. 1d). In contrast to the transcriptional diversity, the performed whole exome sequencing analysis only showed a small number of recurrent genetic alterations. However, since this study aims to integrate a diverse molecular landscape, we included a clinically relevant distribution of the oncogenic KRAS mutations G12D (N=3), G12V (N=3), G12C (N=1), and G12R (N=1) combined with the inclusion of other alterations such as CDKN2A, TP53 and EGFR (Fig. 1c).

Apart from the morphological and molecular variation in PDAC patients, it is crucial to capture the inter-patient treatment response differences. Therefore, both neoadjuvant-treated (N=4) and treatment naïve (N=4) patients were included in this study, where each patient exhibited a distinct clinical response. The clinical response was also recorded as a good, bad, or mixed response, in order to evaluate *ex vitro* organoid response to therapies with matched patient PFS. The representative treatment overview highlights the comprehensive diversity in terms of provided therapy and clinical response (Fig. 1d). Altogether, we have selected and comprehensively characterized a highly diverse panel of 8 PDAC organoid lines which will provide the foundation for the next steps in this study.

### NDR-based intra-well normalization improves patient-stratification over traditional readouts

The aforementioned findings clearly highlight the inter-patient heterogeneity in PDAC, but it is still necessary to determine whether these phenotypic, functional, and molecular differences translate into clinically-relevant *ex vivo* treatment responses. Therefore, we performed a drug screen with the standard of care treatments, using our developed artificial intelligence (AI)-driven image analysis software platform (Orbits). Using computer vision to identify and track organoids, Orbits combines label-free viability assessment with a fluorescent cell death readout. While our clinical data indicate the presence of sensitive and resistant patients to the standard of care treatments, this inter-patient response heterogeneity could not be captured using the standard relative viability readouts of ATP based-assays (Fig. 2a). This is partially due to the high diversity in growth rate within our panel, as it has been reported that fast growing tumors exhibited different therapeutic responses compared to slow growing malignancies^24^. Therefore, we hypothesize that the integration of the individual organoid growth rates into the analysis would improve patient stratification.

**Fig. 2.**
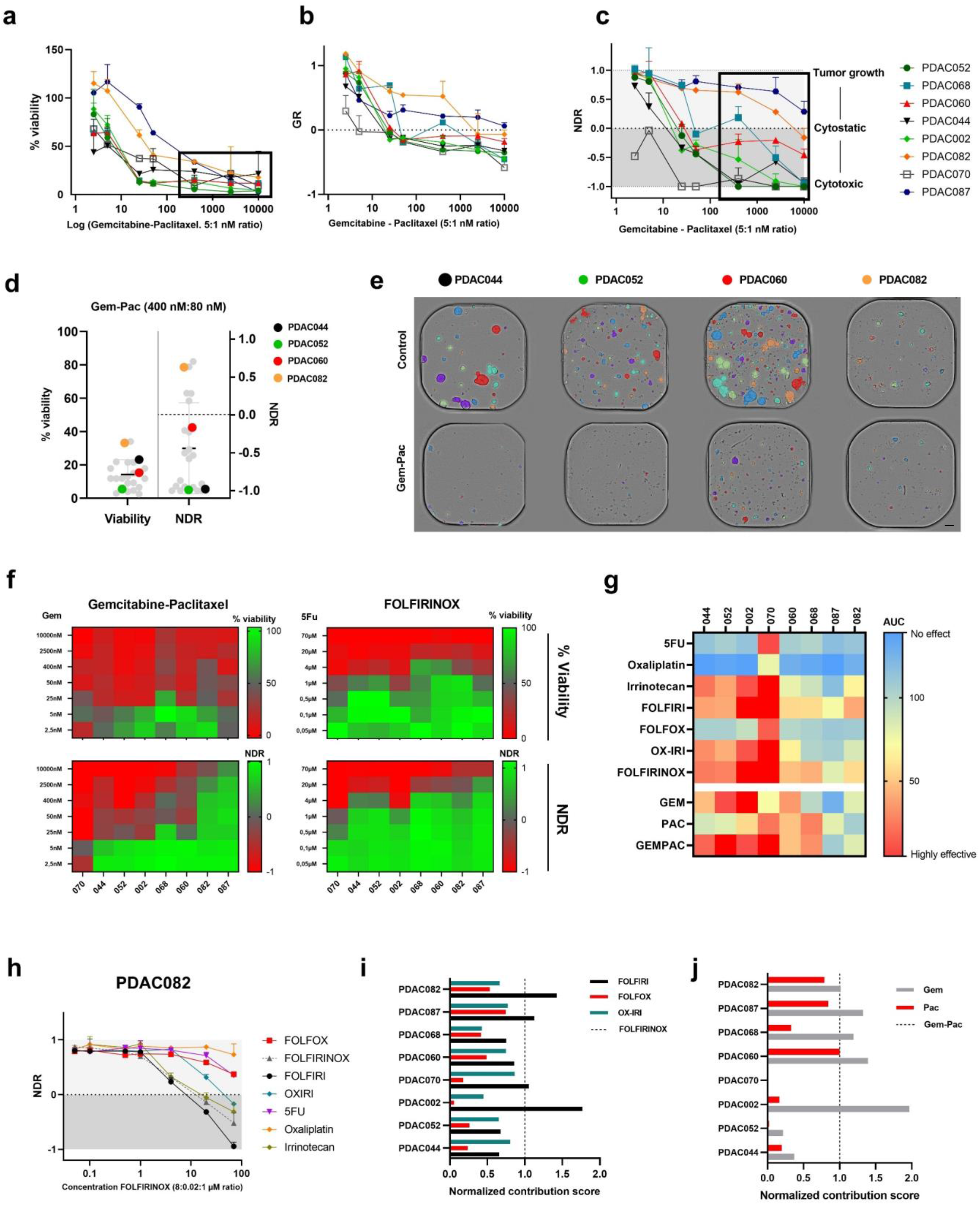
Normalized drug metric (NDR) highlights the inter-patient response heterogeneity. **a-c**. Comparison of relative viability, Growth rate (GR) and NDR (black squares indicate clinically relevant concentration ranges; 400-10000 nM gemcitabine at 5:1 ratio with paclitaxel). **d**. Dot plot comparison of 4 selected PDAC organoid lines (treated with the abovementioned concentration range) showing the increased stratification performance of the NDR. Grey dots represent the other 4 patients. **e**. Label-free annotated brightfield images of PDAC044, PDAC052, PDAC060 and PDAC082 (Orbits software) indicating clear differences in growth (control) and therapy response (400 nM gemcitabine: 80 nM paclitaxel). Scale bar=100 μm **f**. Heatmap-based comparison of % viability and NDR values for gemcitabine-paclitaxel and FOLRIRINOX **g**. NDR-based area under the curve (AUC) indicating patient and therapy specific sensitivities. **h**. NDR-curve of PDAC082 upon treatment with the FOLFIRINOX-based mono and combination treatment. **i-j**. Normalized contribution score (NDR-based) calculation reveals potential patient-specific antagonistic effects of the full combination treatment (FOLFIRINOX and gemcitabine-paclitaxel).

Here, we compared the performance of two growth rate-based normalization metrics: Growth Rate normalization (GR) and Normalized Drug Response (NDR). Interestingly, the intra-well growth rate normalization (GR metric) was not sufficient for discriminating individual patient responses (Fig. 2b) Conversely, the NDR metric, which accounts for variations in seeding density and integrates a positive control (staurosporine), outperforms current response metrics in terms of patient stratification (and the reduction of replicate variation) (Fig. 2c,d, Extended Data Fig. 2a-b). This increased screening performance was also validated on the imaging level, highlighting not only patient-specific growth rates (PDAC060: fast growing and PDAC082: slow growing), but also distinct cytostatic (PDAC060) or cytotoxic (PDAC052) responses to gemcitabine-paclitaxel treatment, which were captured by the NDR metric (both endpoint and kinetic) and not with the % viability readout (Fig. 2d-f, Extended Data Fig. 2c).

As we focus on identifying patient and therapy-specific sensitivities, we treated each PDAC organoid line with a clinically relevant concentration range (taking into account the molar ratios/or intratumoral concentrations without exceeding the peak plasma concentrations) of FOLFIRINOX (5-Fluorouracil; 5-FU:8 μM, irinotecan:0.02 μM, and oxaliplatin:1 μM ratio), gemcitabine-paclitaxel (5 nM:1 nM ratio) and the corresponding mono/modified therapies. In-line with the clinical observations, our NDR-based area under the curve (AUC) analysis revealed patient-specific sensitivities to both the individual and combination therapies (Fig. 2g). Moreover, the NDR metric was also able to capture patient- and therapy-specific antagonistic (oxaliplatin for PDAC082) or additive (gem-pac for PDAC052) effects, which were visualized using the normalized contribution score (ratio AUC of full combination / NDR AUC single therapy) (Fig. 2h-j, Extended Data Fig. 2d-f). In summary, these findings emphasize that neglecting the patient-specific growth rates and intra-well variation (seeding density and organoid size) could confound the observed treatment efficacy via under- or overestimation. Consequently, the use of the standardized NDR metric substantially increased (high-throughput) drug screening performance, leading to better patient-stratification.

### Transcriptomic signatures align with the observed *ex vivo* organoid responses to therapy

While NDR normalization resulted in greater patient stratification *ex vivo*, it is crucial to determine how the measured organoid treatment response reflects the clinical situation. Therefore, we aimed to compare baseline transcriptome signatures for all PDAC organoid lines (N=8) with the NDR metrics from *ex vivo* drug screening. For this, we first grouped sensitive and resistant *ex vivo* responders by conducting a principal component analysis (PCA) (integrating the NDR-based AUC for the standard of care regimen, growth rate, and NDR value at the physiological relevant drug concentration). Based on this analysis, we were able to identify 4 sensitive (PDAC002, PDAC044, PDAC052 and PDAC070) and 4 resistant (PDAC060, PDAC068, PDAC082 and PDCA087) organoid lines (Extended Data Fig. 3a,b). Of note, we also performed a PCA analysis on the normalized read counts (expression level), and we observed a similar clustering of sensitive and resistant patients (Extended Data Fig. 3c). Hierarchical clustering of each subgroup and functional annotation of four gene clusters revealed that the sensitive PDAC organoid cohort had an enrichment in hallmark gene sets such as interferon-alpha response, PI3K-AKT-mTOR signaling, mitotic spindle, apical surface, and epithelial to mesenchymal transition (EMT). On the other hand, the resistant organoid lines showed an increased activation of gene sets related to the fatty acid synthesis, cholesterol homeostasis, IL-6 Jak STAT3 signaling, pancreas β cells and TNFα signaling, which could serve as a rationale for developing more effective treatment strategies (Fig. 3a).

**Fig. 3.**
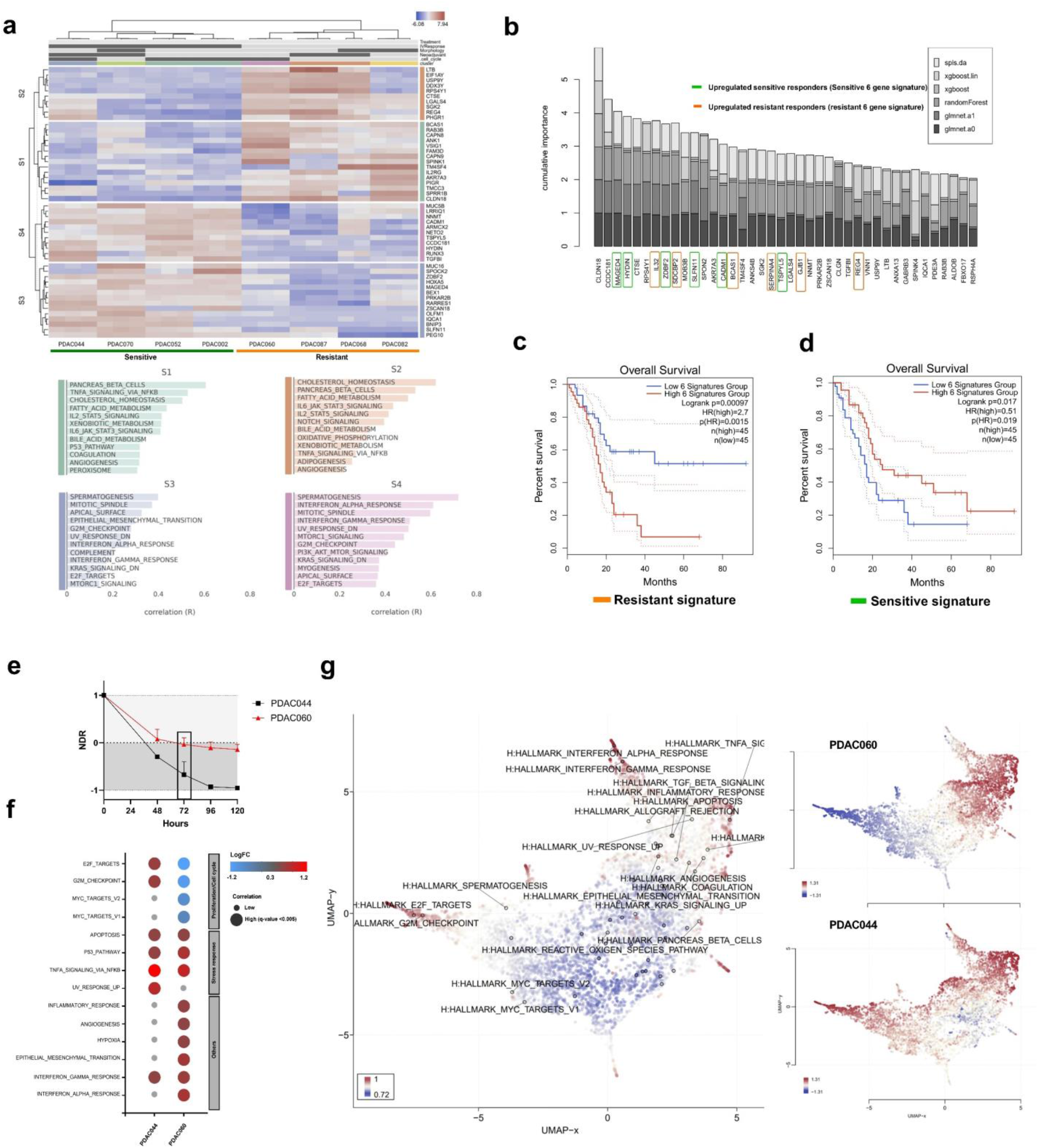
Transcriptional analysis highlights clinical relevance of NDR readout. **a**. Heatmap showing gene expression profiles of good (PDAC044, PDAC070, PDAC052 and PDAC002) and bad (PDAC060, PDAC068, PDAC087 and PDAC082) *ex vivo* responders with the corresponding enriched Hallmark gene set clusters (S1,S2,S3 and S4), sorted by 2-way hierarchical clustering. **b**. Cumulative importance plot, generated by calculating an importance score for each variable (gene) using machine learning algorithms, including LASSO, elastic nets, random forests, and extreme gradient boosting. (Green=upregulated in good responders and orange=upregulated in bad responders). **c-d**. Kaplan-Meier analysis of overall survival using the predefined gene signatures (Good and Bad) using TGCA data. **e**. Kinetic NDR analysis of PDAC044 (cytotoxic responder) and PDAC060 (cytostatic responder) upon treatment with gemcitabine-paclitaxel (400 nM:80 nM). **f**. Bubble plot comparing significantly up- or downregulated gene sets after treatment with gemcitabine-paclitaxel (400 nM:80 nM after 72 hours as indicated with the black square). **g**. UMAP visualization of differently expressed hallmark gene sets following treatment with gemcitabine-paclitaxel (400 nM:80 nM) after 72 hours.

The top 40 differently expressed genes were identified based on the variable importance score to assess if this transcriptional biomarker selection can be extended towards a greater patient cohort. Based on the importance scores, we selected a 6-gene signature for both sensitive (green) and resistant (orange) organoid lines to interrogate the clinical value of the identified transcriptional profiles of both the sensitive and resistant organoid lines in the TCGA PDAC cohort (Fig. 3b). Interestingly, patients with a high expression of the resistant response gene signature had a significantly worse overall survival (OS) (HR=2.7, p=00015), while patients with high expression of the sensitive response gene signature showed a better OS (HR=0.51, p=0.019). Together, these data highlight the value of using the NDR-based drug screening approach, for potential predictive applications in the clinic (Fig. 3c,d, Extended Data Fig. 3d).

In addition to having high clinical relevance, we also assessed whether the image-based drug responses (e.g. cytostatic or cytotoxic) from the NDR-based approach reflect an associated transcriptional state (growth arrest of cell death respectively). As proof-of-concept, we selected PDAC044 and PDAC060, given their specific classification as a cytotoxic (NDR < 0) and cytostatic (NDR = 0) organoid line, respectively (Fig. 3e). Although apoptotic signatures were detected in both patients, PDAC060 showed a substantial decrease in gene sets related to proliferation and cell cycle progression (e.g. E2F targets, G2M Checkpoint, MYC targets V1 and V2), indicating a transcriptional growth arrest state. On the other hand, PDAC044 failed to suppress cell cycle progression and was enriched in gene sets related to cellular stress and apoptosis (e.g. UV response, TNFα Signaling Via NFKβ, apoptosis) (Fig. 3f,g, Extended Data Fig. 3e). Therefore, taken together, the NDR-based approach is not only able to better stratify patient responses, but can also distinguish cytostatic from cytotoxic responses with transcriptomic translatability.

### Single-organoid analysis reveals the intra-tumoral response heterogeneity to therapies

While the NDR metric provided compelling information regarding the mechanism of drug response and increases patient stratification, it still relied on a whole-well (bulk) readout, thereby overlooking intratumoral heterogeneity. Moreover, our clinical data clearly showed that almost all patients (7/8) eventually developed disease progression, implying the presence of treatment-resistant tumor clones (Fig. 4a). Therefore, we aimed to encapsulate the information regarding intratumoral heterogeneity within our PDAC organoid cohort, using our Orbits software analysis platform. Intriguingly, despite having distinct NDR metrics, we identified sensitive and resistant organoid clones in both sensitive (PDAC052) and resistant (PDAC060 and PDAC082) organoid lines upon *ex vivo* treatment with the standard of care therapy, gemcitabine-paclitaxel (Fig. 4b,c). Considering that not all PDAC organoid subclones visually showed the same response to a specific therapy, we developed a single-organoid readout using our Orbits software platform, to quantify individual sensitivities on a high-throughput scale. For this, we used positive and negative controls to first defined the fraction or organoids affected within a well (i.e. cell death) and defined 3 response ranges: resistant (<0.15), sensitive (0.16-0.33), and highly sensitive (>0.34). Our single-organoid endpoint analysis, 5 days after treatment, confirmed visual assessment that resistant clones were present in both sensitive and resistant PDAC organoid lines (Fig. 4d, Extended Data Fig. 4a,b). However, the size and fraction of these resistant PDAC clones clearly varied between the individual patients and does not necessarily line up with the NDR-based sensitivities. For example, PDAC002 had a good NDR-based *ex vivo* response, but the single-organoid analysis detected a vast number of resistant organoid clones (Fig. 4e). Together, this highlights the need for incorporating a single-organoid readout in a predictive clinical setting.

**Fig. 4.**
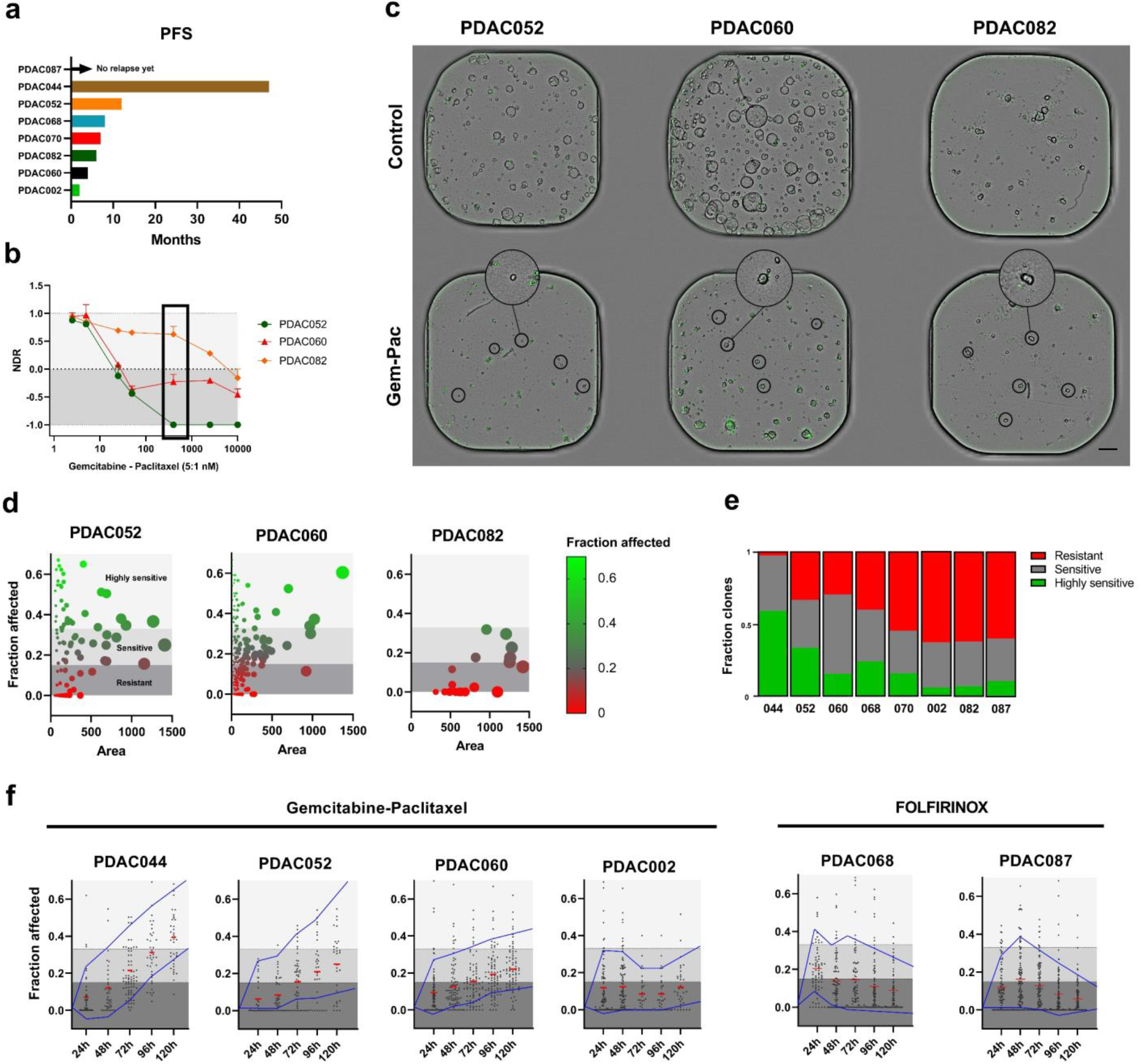
Single-organoid analysis reveals the intra-tumoral response heterogeneity. **a**. Bar plot representation of the progression-free survival (PFS) in months per patient. **b**. NDR results upon treatment with gemcitabine-Paclitaxel highlighting the 3 distinct responses. **c**. Representative brightfield/fluorescence (Cytotox Green) images of PDAC052, PDAC060 and PDAC082 indicating the presence of persistent PDAC organoid clones (black circle). Scale bar=100 μm **d**. Single organoid dose response based on cell death (green area/brightfield area) labeled as fraction affected and area (brightfield) of PDAC052, PDAC060 and PDAC082 treated with gemcitabine-paclitaxel (400 nM:80 nM). Dark grey region (Fraction affected <0.15) indicates resistant, middle grey region sensitive (Fraction affected 0.15-0.34) and light grey highly sensitive (Fraction affected >0.34) PDAC organoid clones. Bubble size correlates with the organoid area. **e**. Relative fraction of resistant, sensitive and highly sensitive PDAC organoids for each patient treated with gemcitabine-paclitaxel (400 nM:80 nM). **f**. Dynamic quantification of single-organoid responses treated with gemcitabine-paclitaxel (400 nM:80 nM) or FOLFIRINOX (20 μM 5-FU:0.0625 μM SN38:2.5 μM Oxaliplatin).

One of the limitations of single-timepoint analysis is that it only provides insight into an isolated window of analysis, and therefore, using time-lapse microscopy, we evaluated single-organoid treatment dynamics over time. This again revealed patient-specific responses, where PDAC044 and PDAC052 showed a clear reduction of resistant clones over time, which is in-line with longer patient PFS (Fig. 4a). By contrast, PDAC060, PDAC002, PDAC068 and PDAC087 still showed a substantial amount of non-responsive organoid clones upon treatment with relatively high concentrations of gemcitabine-paclitaxel (400 nM gemcitabine: 80 nM paclitaxel) or FOLFIRINOX (20 μM 5-FU, 0.0625 μM SN38, and 2.5 μM oxaliplatin) (Fig. 4f, Extended Data Fig. 4c). Moreover, this was also linked to their poor clinical outcome and lower PFS. Overall, this novel readout confirms the patient-specific, intratumoral response heterogeneity in PDAC and subsequently allows us to study this inherent disease promoting feature in more detail.

### Bulk and single-organoid analysis reveals patient-, therapy-, concentration-, and time-specific invasion patterns

In addition to the intratumoral response heterogeneity, PDAC is also characterized by its explicit invasive behavior, resulting in metastatic disease in the majority of patients over time^25-27^. These invasive features are on one hand, intrinsically imprinted or can be promoted by external stress factors such as chemotherapy^28,29^. Based on our live-cell image analysis, we observed strong visual clues (evasion of spindle shaped cells and budding cell clusters) that gemcitabine-paclitaxel (and to a lesser extent FOLFIRONOX) might drive the invasive behavior of PDAC cells (Extended Data Fig. 5a). Moreover, not all PDAC organoid clones within the treated wells showed this invasive behavior, which also appeared to be time-dependent. This again highlights the importance of detecting clonal heterogeneity within organoids to better represent patient tumors in the clinic.

To study these findings more in-depth, we added specific features to our image analysis, enabling us to monitor the invasive regions surrounding the organoids. Quantification was performed by using the sum invasive area, invasive area per organoid area, and invasive fraction (Fig. 5a). By integrating a non-linear dimensionality reduction algorithm (tSNE), we were able to confirm our finding (tested as proof-of-concept on an invasive patient; PDAC002) that not all PDAC organoid clones are driven in the same ‘direction’ (no uniform response) following treatment with gemcitabine, paclitaxel or the combination at a clinically relevant concentration range. The tSNE analysis revealed the presence of distinct clusters, highlighting the response heterogeneity. In this context, cluster 6 represents PDAC organoids with invasive characteristics (high invasive area and high invasive fraction), cluster 7 sensitive PDAC organoids (high fraction affected) and cluster 2 and 5 PDAC organoids with an increased area and lower fraction affected (healthy, growing organoids) (Fig. 5b, Extended Data Fig. 5b-e). The conducted pseudotime analysis (trajectory interference analysis) also indicated temporal and treatment dependent transition trajectories towards the invasive, sensitive and non-responsive PDAC organoids (Fig. 5c). Interestingly, we observed a strong overlap (close proximity on tSNE plot) between the invasive (cluster 6) and sensitive clusters (cluster 7), suggesting the presence of both sensitive and less-responsive organoid clones with invasive features (Fig. 5d). Furthermore, to assess whether these invasive features are treatment specific, we calculated the distribution (validated with the paclitaxel *ex vivo* response, Extended Data Fig. 5f) over the invasive trajectory for gemcitabine, paclitaxel or the combination regimen. As indicated on the imaging level, paclitaxel shows the highest organoid density over this path, confirming the initial indications that indeed chemotherapy (paclitaxel) might promote the invasive characteristics of PDAC (Fig. 5e).

**Fig. 5.**
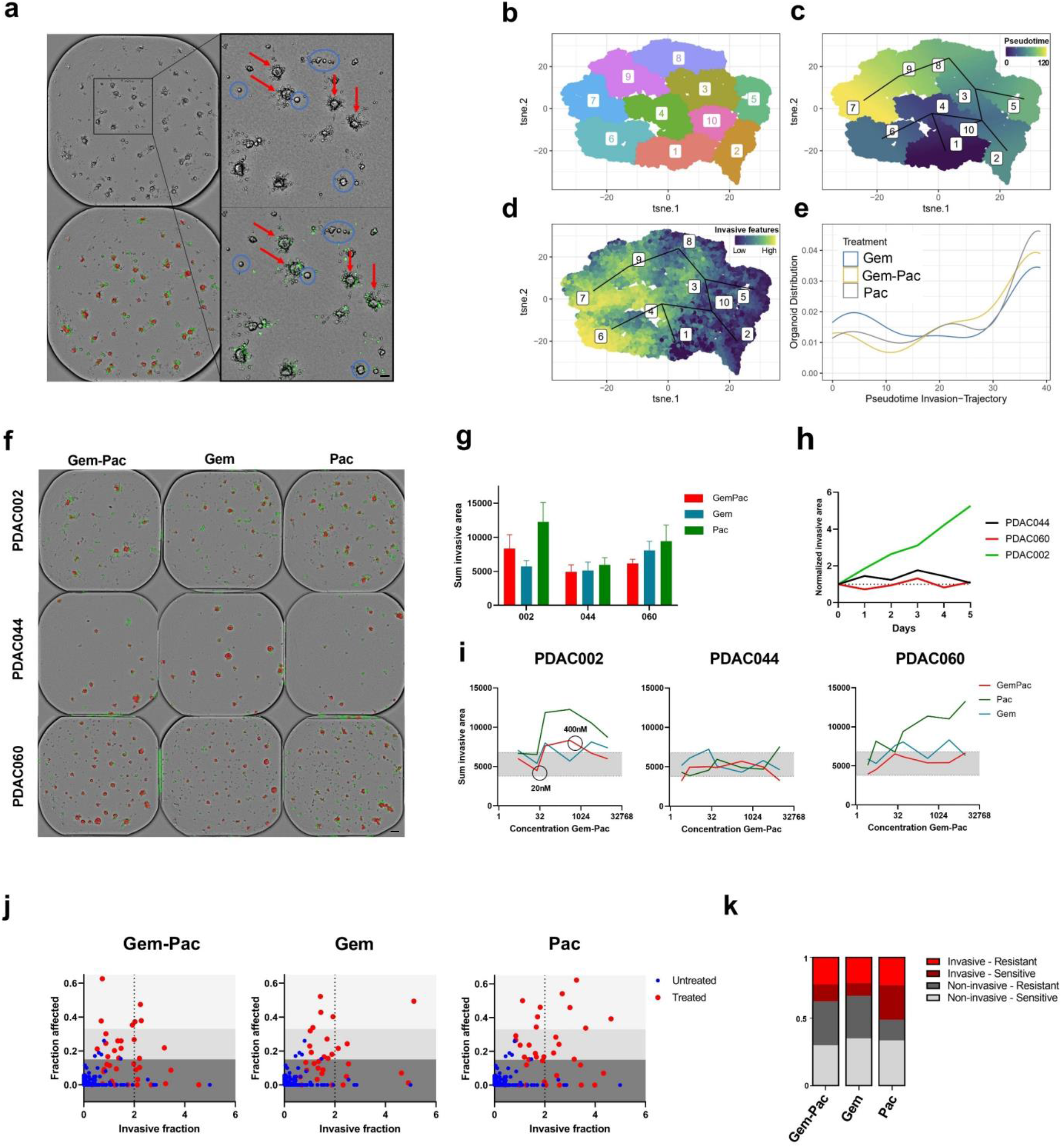
PDAC organoids reveal patient-, therapy-, concentration- and time-specific invasive patterns. **a**. Representative brightfield (top), masked (left under) and fluorescent overlap (right under) images of PDAC002 highlighting the clone specific invasive behavior. Red arrows= invasive PDAC organoid clones, blue circles=non-invasive PDAC organoid clones. Scale bar=50 μm. **b**. tSNE analysis of 6 features (area, invasive area, fraction affected, invasive fraction, brightness and texture) resulting in 10 distinct clusters of single PDAC002 organoids (untreated or treated with gemcitabine, paclitaxel and gemcitabine-paclitaxel). **c**. Whole 384-well plate (all conditions) single-organoid pseudotemporal trajectory analysis of patient PDAC002. **d**. tSNE visualization of PDAC organoids with a high (yellow) invasive fraction. **e**. Therapy comparison of the PDAC density distribution over the invasive trajectory (cluster 6). **f**. Annotated brightfield images masking the PDAC organoids (red) and invasive area (green) upon treatment with gemcitabine-paclitaxel (400 nM:80 nM) and the corresponding monotherapies. Scale bar=100 μm **g**. Endpoint quantification of the sum invasive area. **h**. Kinetic quantification (5 days) of the sum invasive area upon treatment with gemcitabine-paclitaxel normalized to timepoint 0. **i**. Concentration dependent invasive behavior of PDAC002, PDAC044 and PDAC060. **j**. Single-organoid endpoint analysis of PDAC002 treated with either Gem-Pac (400 nM:80 nM), Gem (400 nM) and Pac (80 nM). **k**. Relative quantification of invasive, non-invasive, sensitive and resistant PDAC organoid clones of PDAC002. Gem-Pac=gemcitabine-paclitaxel (400 nM: 80 nM), Gem=gemcitabine (400 nM) and Pac=paclitaxel (80 nM).

Next, we investigated whether the developed organoid analysis can accurately quantify patient- and concentration specific differences in terms of invasiveness over time. As seen visually, PDAC044 showed a minimally invasive area upon treatment, whereas paclitaxel clearly increased the invasive area around the organoids of PDAC060 and PDAC002 (Fig. 5f). However, kinetic and endpoint analysis revealed that the addition of gemcitabine drastically reduced this invasive behavior of PDAC060, which was not observed for PDAC002, again emphasizing patient-specific behavior (Fig. 5g,h). Moreover, these therapy-specific invasive characteristics also appeared to be concentration dependent, highlighting the complexity of this disease-promoting phenomenon (patient-therapy-concentration specific) (Fig. 5i).

Lastly, considering that the tSNE analysis indicates the presence of sensitive and resistant invasive organoid clones, we evaluated the invasive behavior on single-organoid resolution. In-line with the density analysis, paclitaxel shows the highest increase in both sensitive and resistant invasive clones (Fig. 5j,k). When translating this to the clinical response, it is of interest that the addition of paclitaxel to the initial gemcitabine monotherapy increased the clinical response *in vivo* (*ex vivo* observation: increase in sensitive clones with paclitaxel and the combination therapy). However, eventually, the patient (PDAC002) still developed liver metastasis and disease progression due to the presence of resistant clones (*ex vivo* observation: high number of invasive-resistant clones with the combination treatment). Overall, these findings emphasize the extensive intra-tumoral heterogeneity within the invasive landscape, which is orchestrated on a concentration, therapy, patient and time specific level. Moreover, this data also suggests further exploration on how chemotherapy (paclitaxel) can paradoxically fuel the invasive characteristics of PDAC.

### Combination of the NDR metric with a single organoid analysis improves the clinical translatability

While patient-derived tumor organoids have shown initial promise for predicting drug sensitivity of individual patient tumors, a recent prospective intervention trial (SENSOR trial) revealed that only a subset of patients benefited from the organoid technology^30^. The limited predictivity in this study was largely due to the limitations of traditional assays that use rudimentary, bulk readouts (e.g. CellTiter-Glo 3D viability assay), which flattens the complexity of patient tumors, thus hampering wide clinical translatability. Considering this unmet need, we evaluated the approach of combining the NDR metric (e.g. type of response) with data on single-organoid level (e.g. presence of resistant subclones), to improve the predictive potential of organoids in the clinic. To study this, we retrospectively investigated the PFS of the individual patients and matched their clinical response to their *ex vivo* organoid response at single-organoid resolutions. Based on the previously established response ranges, we were able to quantify the fraction of resistant, sensitive, and highly sensitive clones for each individual patient. Linear regression analysis for clinically relevant concentrations of FOLFIRINOX or gemcitabine-paclitaxel, showed a significant correlation with either the percent sensitive (R=0.86 and p=0.0023) or percent resistant clones (R=0.64 and p=0.029) (Fig. 6a,b and Extended Data Fig. 6a,b). However, the ratio of sensitive and resistant clones demonstrated the most superior predictive ability of PFS (R=0.97 and p=<0.0001) and therapy response (Fig. 6c,d).

**Fig. 6.**
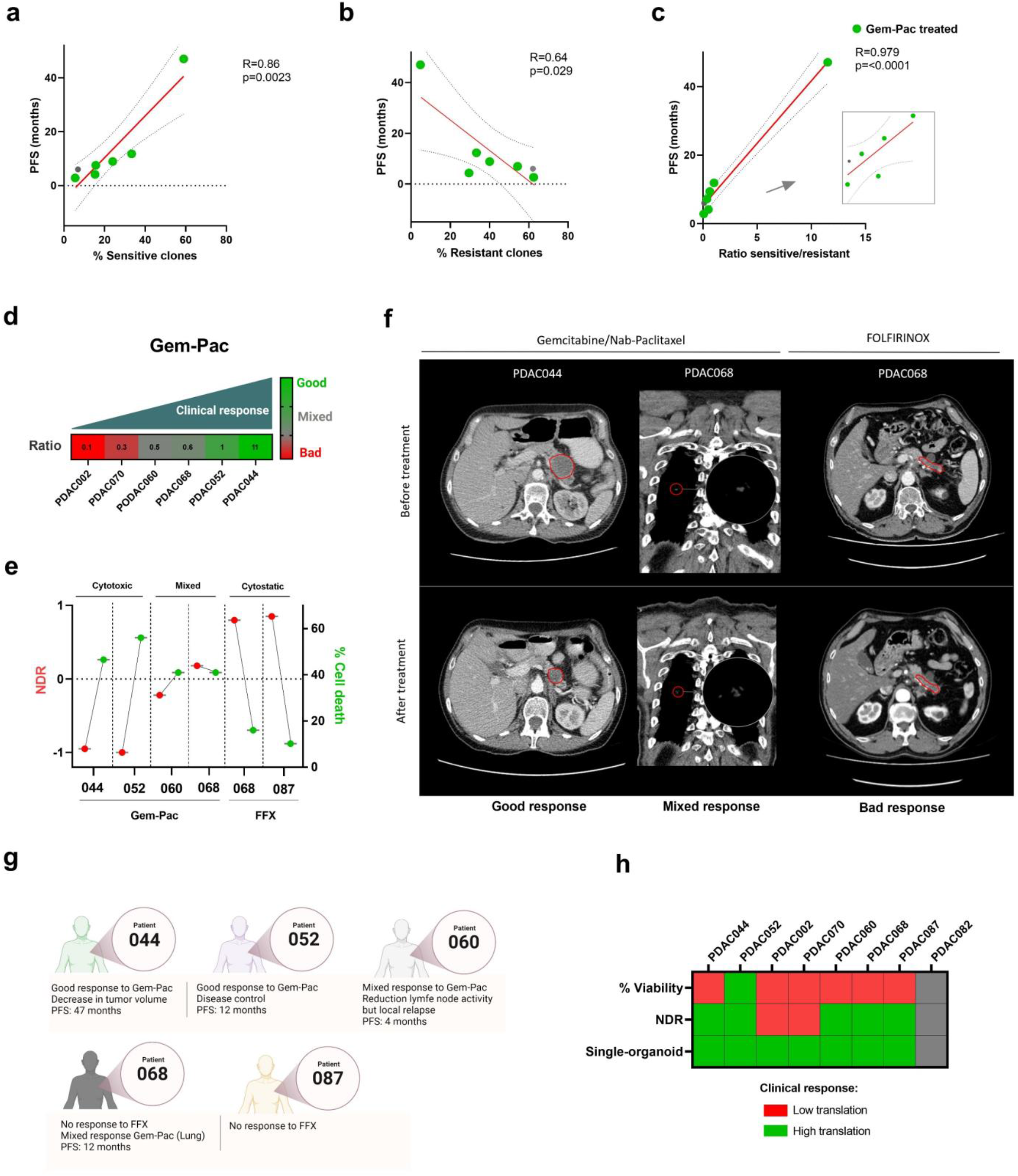
Clinical significance of NDR and single organoid analysis. **a-c**. Correlation of the % sensitive, % resistant and ratio % sensitive/resistant PDAC clones (gemcitabine-paclitaxel; 400 nM: 80 nM) with the PFS. **d**. Correlation of % ratio sensitive/resistant with the initial clinical response to gemcitabine-paclitaxel. Responses were based on available radiological CT protocols. Good= tumor regression and/or PFS >10 months, Mixed= minor tumor regression, Bad= no tumor regression and/or fast disease progression. **e**. NDR signatures indicating the NDR value (left y-axis) and % cell death (right y-axis) upon treatment with gemcitabine-paclitaxel (400 nM:80 nM) or FOLFIRINOX (4 μM 5-FU:0.0125 μM SN38:0.5 μM Oxaliplatin). **f**. Representative CT-scans before and after treatment with gemcitabine/nab-paclitaxel or FOLFIRINOX (8 cycles). Red contours indicate the tumor margins. **g**. Overview of the clinical characteristics of patient PDAC044, PDAC052, PDAC060, PDAC068 and PDAC087. **h**. Comparison of the implemented readouts, suggesting a potential clinical application of combining the NDR readout with the developed single-organoid analysis.

Nevertheless, despite the ability of our developed the single organoid analysis to identify treatment resistant and sensitive organoid subclones, it currently neglects the type of response. Therefore, in addition to correlating the presence of resistant subclones with the PFS, we also aimed to evaluate whether specific NDR signatures can predict the type of response to the standard of care therapies. For this, we combined the NDR metric with the % cell death in order to develop the following response signatures: cytotoxic (NDR < -0.5 and cell death > 40%), mixed response (cell death detectable; 0 < NDR > -0.5 and cell death >40%) and cytostatic (minimal cell death detectable; NDR > 0 and cell death < 40%) (Fig. 6e). Based on these thresholds, PDAC044 showed a cytotoxic signature under gemcitabine-paclitaxel treatment, which correlated with a regression of the primary tumor (−27%) and a reduction of the metastatic liver nodules (not shown in the provided z-plane) (Fig. 6f). On the other hand, PDAC068 and PDAC060 were characterized by a mixed clinical response under gemcitabine-paclitaxel treatment (PDAC068: slight regression of metastatic lung nodules, PDAC060: regression of a lymph node metastasis but only disease control of the primary tumor site), which was in-line with the mixed response signature observed *ex vivo* (cytostatic NDR around 0 but still a high amount of cell death).

We also evaluated patients with no tumor regression under the FOLFIRINOX regimen (PDAC068 and PDAC87). Interestingly, these patients were characterized by a cytostatic signature, again matching the clinical outcome (no tumor regression under FOLFIRINOX treatment (Fig. 6f,g, Extended Data Fig. 6c).

Finally, when evaluating the clinical translatability of all included readouts, the current gold standard (% viability) showed almost no trend with the clinical outcome (e.g. PDAC044 was more resistant then PDAC060, while the clinical data showed the opposite). On the other hand, the NDR signatures (bulk readout) correlated with the initial clinical response in a subset of patients (5/8, 62%) (Fig. 6h). However, the addition of the single organoid analysis showed a high translatability in all patients, emphasizing the predictive power of combining these 2 measurements in one single readout, which is capable of predicting the response to therapy as well as the probability of developing relapse due to the presence of resistant PDAC clones (Fig. 6h).

## Discussion

Over the past years, tumor organoids have been widely adopted in both preclinical (development of novel drug candidates) and clinical research (personalized medicine)^30,31^. Nevertheless, despite their numerous advantages and increased complexity, we still rely on primitive viability readouts that have been used for decades in traditional two-dimensional cell cultures. As a result, we completely neglect the response heterogeneity of individual PDAC clones, thereby diminishing the translational relevance. Considering that PDAC is characterized by a high relapse rate due to the presence of invasive and treatment resistant tumor cells, it is essential that more advanced assays are incorporated into the rapidly evolving field of “patients-in-a-lab”. For this study, we developed a novel automated high-throughput screening pipeline that can dynamically study the type of response as well as the subclonal response heterogeneity on single-organoid resolution. In accordance with the variance in molecular landscape and clinical response, our *ex vivo* chemotherapy screen revealed patient-specific treatment sensitivities. Importantly, these distinct response patterns were only evaluable with the NDR metric and not with the GR and relative viability readout. Furthermore, as explained above, an important aspect of this chemotherapy screen was to use clinically relevant concentrations/combinations of the individual monotherapies^32-34^. In-line with previous findings, we show that oxaliplatin monotherapy shows limited efficacy, whereas irinotecan and 5-FU drive the efficacy of the FOLFIRONOX regimen^35,36^. However, using the NDR readout, we were able to capture patient-specific additive effects of oxaliplatin in combination with 5-FU, confirming the rationale for combining all three chemotherapeutics. With respect to the NDR metric, it needs to be taken into account that for slow growing tumor organoids (growth rate <1.2), the NDR might overestimate the initial effect (e.g. PDAC070). Nonetheless, even though the NDR shows an overall better drug screening performance (can distinct cytostatic from cytotoxic responses and shows conformity with the transcriptional signatures) compared to the current gold standards, we still did not capture the full story of PDAC as heterogenous and immensely complex/adaptive disease^37,38^. Therefore, we integrated a novel single-organoid readout that is able to dynamically monitor this complexity and response heterogeneity. Interestingly, as documented in pathological tumor regression assessments, we were able to identify persistent malignant ducts, even with patients that showed a good NDR-based *ex vivo* response^39^. This presence of less sensitive clones (even at clinically relevant concentrations) could explain why eventually 7 out of 8 patients had disease progression due to local or metastatic outgrowth of the tumor.

Furthermore, in accordance with the high metastatic burden in PDAC, it is well described in previous studies that chemotherapy-induced stress might promote invasion and EMT^40-43^. To our interest, we also observed time, concentration, therapy, and patient-specific differences in invasive cells/cell clusters that migrate out of the PDAC organoids. However, a limitation of this study is that we not functionally confirmed the mesenchymal/invasive state of these cells due to the technical difficulty of specifically isolating this small subset of cells. Nevertheless, morphological comparison of our organoids with published organoid invasion studies combined with the identification of spindle shape cells/budding cell clusters migrating out of the organoid upon treatment suggest an EMT-like state (which can be organoid clone specific)^44-47^. Moreover, as indicated by the tSNE analysis, we observed a vast number of invasive-resistant PDAC organoid clones upon treatment with gemcitabine-paclitaxel, which might explain the course of PDAC as a highly invasive and treatment resistant disease over time.

In the end, when evaluating the clinical relevance of our findings, it should be addressed that this study serves as a proof-of-concept, since we only include 8 patients diagnosed with PDAC. Therefore, we highly anticipate on evaluating the clinical relevance of our analysis pipeline in a greater cohort of patients. To support the (clinical) validation of our approach by other groups, a detailed protocol has been published^22^ and our Orbits image analysis platform will be made available in a cloud-based platform. Our data, once again, highlights the inter-patient and therapy-specific response heterogeneity in PDAC, and our organoid analysis approach can detect these differences. This will become a critical tool for overcoming tumor and patient heterogeneity in personalized medicine.

In addition to the clinical applications, this innovative readout can also serve as a foundation for developing novel/more effective treatments for PDAC (and other tumor types). Especially because it will allow researchers to evaluate the efficacy of a novel therapeutic on multiple subclones (i.e. does the therapy targets all subclones), it will provide them information regarding potential pro-invasive properties (i.e., does the therapy increase the invasiveness) and it is fully automated/high-throughput compatible. To conclude, we developed and validated a novel AI-driven live-cell imaging readout that enabled us to study the complexity and intrinsic subclonal response heterogeneity and invasive properties of PDAC in more depth. Consequently, we are convinced that these findings will inevitably breathe new life into the rapidly evolving landscape of personalized medicine for which patient-derived tumor organoids will inevitably become an important predictive platform.

## 3. Methods

### Establishment of human pancreatic cancer organoids

Tissue resection fragments were obtained from PDAC patients undergoing curative surgery at the Antwerp University Hospital. Written informed consent was obtained from all patients, and the study was approved by the UZA Ethical Committee (ref. 14/47/480). The human biological material used in this publication was provided by Biobank@UZA (Antwerp, Belgium; ID: BE71030031000); Belgian Virtual Tumorbank funded by the National Cancer Plan. The corresponding resected tumor fragments were stored in Ad-DF+++ (Advanced DMEM/F12 (GIBCO), with 1% GlutaMAX (GIBCO), 1% HEPES (GIBCO), 1% penicillin/streptomycin (GIBCO) supplemented with 2% Primocin (Invivogen) at 4°C and transported on ice to be processed within 24 hours for organoid culture. The tumor fragments were dissected on ice to remove remaining connective (healthy) tissue and are subsequently cut into small pieces of approximately 2 mm in diameter. Following dissection, the tissue fragments were washed with ice cold phosphate-buffered saline (PBS) and were subsequently enzymatically digested at 37°C for 1h using 5 mg/mL collagenase II (Sigma-Aldrich), 10 μM Y-27632 (Cayman Chemicals) and 1:500 Primocin (Invivogen). After digestion, the cell suspension was passed through a 100 μm filter (Fischer Scientific) and the filtrates were centrifuged at 300 x g for 5 min. The strained cell pellet was resuspended in >80% ice cold Cultrex growth factor reduced BME type 2 (R&D Systems) in PDAC organoid medium. Small droplets of 20 μL were plated and were incubated inverted for 30 minutes at 37°C to allow them to solidify. Thereafter, the droplets were overlayed with Full PDAC medium consisted of 0.5 nM WNT Surrogate-Fc-Fusion protein (ImmunoPrecise), 4% Noggin-Fc Fusion Protein conditioned medium (ImmunoPrecise), 4% Rspo3-Fc Fusion Protein conditioned medium (ImmunoPrecise), 1x B27 without vitamin A (Gibco), 10 mM nicotinamide (Sigma-Aldrich), 1 mM N-acetylcysteine (Sigma-Aldrich), 100 ng/ml FGF-10 (Peprotech), 500 nM A83-01 (Tocris), 10 nM gastrin (R&D Systems) and 10 μM Y-27632 (Cayman Chemicals). To prevent microbial contamination, 1x Primocin was added to the culture medium during the first two weeks. For passaging, the organoids were digested to single cells with TrypLE Express (GIBCO). For cryopreservation, 3-day old organoids were harvested with Cultrex Harvesting Solution (R&D Systems) and frozen in Recovery Cell Culture Freezing Medium (GIBCO). Samples were tested for Mycoplasma contamination with the MycoAlert Mycoplasma Detection Kit (LONZA).

### Histological evaluation

Early passage organoids were collected using Cultrex Organoid Harvesting Solution, washed with ice-cold PBS, and fixated in 4% paraformaldehyde for 30 minutes at room temperature. Fixed organoids were transferred to a 4% agarose micro-array mold and paraffin embedded as described before^48^. Five μm-thick sections were prepared, deparaffinized and rehydrated prior to Hematoxylin+Eosin (H&E) staining. Images were acquired on the Leica DM750.

### Drug screening

Drug screening was performed using our pre-validated drug screening pipeline for which a detailed protocol is available in the Journal of Visualized Experiment^49^. Briefly, three days before the start of the experiment, the PDAC organoids were passaged as single cells using TrypLE and plated in Cultrex drops. Subsequently, the PDAC organoids were harvested (enzymatic digestion) with Cultrex Harvesting Solution, collected in 15 mL tubes coated with 0.1% BSA/PBS, washed with Ad DF+++ and resuspended in 1 mL Full Ad-DF+++ medium (without Y-27632). Next the number of organoids were counted with the in-house developed organoid counting software in a 384-well microplate (Orbits). The PDAC Organoids were then diluted in Full Ad-DF+++ and 4% Cultrex on ice to a concentration that results in ∼200 organoids/50μL. Next, 50μL of this solution was dispensed into a 384-well ultra-low attachment microplate (Corning, #4588) using the OT-2 a pipetting robot (Opentrons) in a cooled environment. Thereafter, the plate was centrifuged (100 rcf, 30 sec, 4°C) and incubated overnight at 37°C allowing the organoids to recover from the stress during plating.

All drugs and fluorescent reagents were added to the plate using the Tecan D300e Digital Dispenser. Cytotox Green (60 nM / well, Sartorius), Staurosporine (2 μM), 5-Fluorouracil (5-FU), SN38 (active metabolite Irinotecan), gemcitabine and paclitaxel (MedChemExpress) were dissolved in DMSO. Oxaliplatin and leucovorin (MedChemExpress) were dissolved in PBS to yield a final concentration of 0.3 Tween-20 required for dispensing with the D300e Dispenser. To mimic the clinical setting, the following ratios were used based on tissue relevant concentration ranges (not exceeding peak plasma concentrations): gemcitabine-paclitaxel (5 nM gemcitabine:1 nM paclitaxel) and FOLFIRONOX (20 μM 5-FU:0.0625 μM SN38:2.5 μM Oxaliplatin). A fixed concentration of Leucovorin (1 μM) was used to enhance the efficacy of 5-FU (as in the clinical setting). Brightfield and green fluorescence whole-well images (4x objective) were taken every 24 hours with the Tecan Spark Cyto set at 37°C / 5% CO2 for 5 days.

### Drug response metrics

Following image acquisition with the Tecan Spark Cyto, Brightfield and fluorescence images were analyzed using the validated and in-house developed analysis software Orbits platform. For % viability assessment, data results were normalized to vehicle (100%) and/or baseline control (0%) (Staurosporine 2 μM). The growth rate (GR) metric (negative control) and the normalized drug response (NDR) metric (positive and negative controls) were calculated based on the work of Hafner and colleagues and Gupta and colleagues, respectively, using the adapted R script^50,51^. The Total brightfield Area – Total Green Area parameter was used, and the fold change was calculated for each well individually from the first measurement (T0) and a timepoint as indicated in the figures. Based on the NDR values, the drug effects can be classified as: >1, proliferative effect; = 1, normal growth as in negative control; = 0, complete growth inhibition; = -1, complete killing.

### Single-organoid analysis

For each organoid, the label free total masked area (Orbits) and overlapping total green area (Cytotox green) was calculated. Based on this ratio (total green/total masked area, fraction affected), we defined the following response ranges: resistant (<0.15), sensitive (0.16-0.33), and highly sensitive (>0.34). For the quantification of the invasive characteristics, we improved our orbits analysis to quantify the survival invasive area (masked invasive are – overlapping green invasive area) and the invasive fraction (invasive area/masked organoid area). An invasive fraction <2 indicates tumor organoids with invasive characteristic. To model temporal heterogeneity in the response profiles of the organoid population obtained from a single patient (i.e. PDAC002), the morphology, treatment response and invasive cancer cell behavior at different time points (i.e. baseline to 5 days) were subjected to principal component analysis (PCA) using the BioC-package *PCAtools*. In total, 29,371 data points were included in the analysis. Measurements were centered and scaled to unit variance. Thousand random matrix permutations (i.e. Holm’s method) were used to select informative principal components (PCs) by comparing the observed and expected percentage of variation explained by each PC. Next, informative PCs were subjected to t-Stochastic Neighbor Embedding (t-SNE) using the R-package *Rtsne*, with the perplexity hyperparameter set to the root square of the number of organoids. Then, the t-SNE output was analyzed using k-nearest neighbor searching (R-package *FNN*) with k set to 1,000, resulting in a network in which each data point is connected to its 1,000 nearest neighbors with connection strengths (i.e. edge weights) set to the complement of the Euclidean distance between the connected samples in the t-SNE output. The resulting weighted network was analyzed using the louvain community detection algorithm (R-package *igraph*) to identify clusters, which were further characterized by mapping measurements of organoid morphology, treatment response and invasive behavior onto the clustered scatter plot. Prior to visualization, data were rank normalized to enhance contrast. Next, using the BioC-package *TSCAN*, pseudotime vectors were calculated based on the t-SNE coordinates and the cluster labels. Therefore, a minimal spanning tree (MST) was first calculated where each node is a cluster centroid, and each edge is weighted by the Euclidean distance between centroids. This represents the most parsimonious explanation for a particular trajectory and has the advantage of being directly interpretable with respect to the pre-existing clusters. The MST was forced to end in clusters of data points representing resistant, sensitive or invasive organoids, as these states are the most discrete in the data set. Then, each data point was mapped onto the closest edge of the MST and pseudotime vectors were computed according to each path through the MST starting from the cluster most strongly enriched for baseline measurements. This was done to ensure that each pseudotime vector recapitulated chronologic changes in real-time. Common pseudotime is defined as the pseudotime averaged across all paths.

### RNA Sequencing

For RNA sequencing (RNA seq), full grown organoids PDAC organoids were harvested from the 384-well plate to mimic the same conditions as the drug screen. Afterwards, RNA was extracted using RNeasy midi kit (Qiagen) for tumor samples of 20-250 mg. For removal of gDNA, RNAse-free DNAse treatment was performed. RNA concentration and purity were checked using the Qubit RNA BR Assay Kit on Qubit 4 Fluorometer (ThermoFisher) and NanoDrop ND-1000 (ThermoFisher), respectively. Samples were frozen at -80 °C and delivered to Genomics Core Leuven for transcriptome sequencing using Lexogen QuantSeq 3’ FWD library preparation kit for Illumina on a Hiseq400 SR50 line with a minimum of 2M reads per sample. Downstream analysis (Differential gene expression, Gene set enrichment, PCA, UMAP and Cumulative importance analysis) was performed using the validated Omics Playground tool of Big Omics Analytics.

### Whole exome sequencing

Full grown PDAC organoids were collected as described above and DNA was isolated using the QIAamp DNA blood mini kit (Qiagen) and send to Genewiz Europe (Leipzig, Germany) for whole exome sequencing on an Illumina NovaSeq platform (2 × 150 bp sequencing, 12Gb (120x)). Raw reads 24 were mapped onto the human reference genome (hg38) using the BWA MEM algorithm (v.0.7.17) in standard settings. Resulting SAM-files were converted into BAM-files, coordinated-sorted, and indexed using samtools (v.1.9). Reads overlapping with 86 genes mutated in at least 10% of either PDAC samples included in the cBioPortal for cancer genomics were selected using the bedtools (v.2.29.2) intersect-command retaining the full read in the original BAM-file (−wa 219 option). Variants were called using the haplotype-based variant detector freebayes (v.1.3.2; 220 Garrison E, Marth G. Haplotype-based variant detection from short-read sequencing. arXiv available under aCC-BY-NC-ND 4.0 International license. (which was not certified by peer review) is the author/funder, who has granted bioRxiv a license to display the preprint in perpetuity. It is made bioRxiv preprint doi: https://doi.org/10.1101/2021.09.09.459656; this version posted October 5, 2022. The copyright holder for this preprint 11 221 preprint arXiv:1207.3907 [q-bio.GN] 2012) in standard settings and indel positions were left aligned and normalized using the bcftools (v.1.9) norm-command. Resulting VCF-files were annotated using SnpEff (v.5.1) for functional annotations and SnpSift (v.5.1) for human genetic variation using dbSNP (build 154), ClinVar (release 05/09/2022) and COSMIC (v96). Annotated VCF files were further manipulated using the SnpSift filter-command retaining only coding, non-synonymous variants that are contained COSMIC, that are not considered as benign and/or likely benign by ClinVar, that have a minimal coverage of 120x with at least 40 reads supporting the alternate allele with no positional or strand bias, that are not of germline origin or are reported as common variants (minor allele frequency>1%) in any human population and that are either missense, stop gained or frame shift mutations. Mutational profiles were visualized in oncoprint format.

## References

1. Mizrahi, J.D., Surana, R., Valle, J.W. & Shroff, R.T. Pancreatic cancer. Lancet 395, 2008–2020 (2020).

2. Monberg, M.E., et al. Occult polyclonality of preclinical pancreatic cancer models drives in vitro evolution. Nat Commun 13, 3652 (2022).

3. Ferrone, C.R., et al. Pancreatic ductal adenocarcinoma: long-term survival does not equal cure. Surgery 152, S43–49 (2012).

4. Sohal, D.P.S., et al. Metastatic Pancreatic Cancer: ASCO Clinical Practice Guideline Update. J Clin Oncol 36, 2545–2556 (2018).

5. Gnanamony, M. & Gondi, C.S. Chemoresistance in pancreatic cancer: Emerging concepts. Oncol Lett 13, 2507–2513 (2017).

6. Seth, S., et al. Pre-existing Functional Heterogeneity of Tumorigenic Compartment as the Origin of Chemoresistance in Pancreatic Tumors. Cell Rep 26, 1518–1532 e1519 (2019).

7. Hwang, W.L., et al. Single-nucleus and spatial transcriptome profiling of pancreatic cancer identifies multicellular dynamics associated with neoadjuvant treatment. Nat Genet (2022).

8. Raghavan, S., et al. Microenvironment drives cell state, plasticity, and drug response in pancreatic cancer. Cell 184, 6119–6137 e6126 (2021).

9. Seppala, T.T., et al. Patient-derived Organoid Pharmacotyping is a Clinically Tractable Strategy for Precision Medicine in Pancreatic Cancer. Ann Surg 272, 427–435 (2020).

10. Hadj Bachir, E., et al. A new pancreatic adenocarcinoma-derived organoid model of acquired chemoresistance to FOLFIRINOX: First insight of the underlying mechanisms. Biol Cell (2021).

11. Yao, J., et al. A pancreas tumor derived organoid study: from drug screen to precision medicine. Cancer Cell Int 21, 398 (2021).

12. Ooft, S.N., et al. Patient-derived organoids can predict response to chemotherapy in metastatic colorectal cancer patients. Sci Transl Med 11(2019).

13. Driehuis, E., et al. Pancreatic cancer organoids recapitulate disease and allow personalized drug screening. Proc Natl Acad Sci U S A (2019).

14. Sachs, N., et al. A Living Biobank of Breast Cancer Organoids Captures Disease Heterogeneity. Cell 172, 373–386 e310 (2018).

15. Kijima, T., et al. Three-Dimensional Organoids Reveal Therapy Resistance of Esophageal and Oropharyngeal Squamous Cell Carcinoma Cells. Cell Mol Gastroenterol Hepatol 7, 73–91 (2019).

16. Phan, N., et al. A simple high-throughput approach identifies actionable drug sensitivities in patient-derived tumor organoids. Commun Biol 2, 78 (2019).

17. Depaoli, M.R., et al. Real-Time Imaging of Mitochondrial ATP Dynamics Reveals the Metabolic Setting of Single Cells. Cell Rep 25, 501–512 e503 (2018).

18. Niepel, M., Hafner, M., Chung, M. & Sorger, P.K. Measuring Cancer Drug Sensitivity and Resistance in Cultured Cells. Curr Protoc Chem Biol 9, 55–74 (2017).

19. Wiley, C.D. & Campisi, J. From Ancient Pathways to Aging Cells-Connecting Metabolism and Cellular Senescence. Cell Metab 23, 1013–1021 (2016).

20. Martins, I., et al. Chemotherapy induces ATP release from tumor cells. Cell Cycle 8, 3723–3728 (2009).

21. Deben, C., et al. OrBITS: label-free and time-lapse monitoring of patient derived organoids for advanced drug screening. Cell Oncol (Dordr) (2022).

22. Le Compte, M., et al. Multiparametric Tumor Organoid Drug Screening Using Widefield Live-Cell Imaging for Bulk and Single-Organoid Analysis. J Vis Exp (2022).

23. Zaid, M., et al. Imaging-Based Subtypes of Pancreatic Ductal Adenocarcinoma Exhibit Differential Growth and Metabolic Patterns in the Pre-Diagnostic Period: Implications for Early Detection. Front Oncol 10, 596931 (2020).

24. Mehrara, E., Forssell-Aronsson, E. & Bernhardt, P. Objective assessment of tumour response to therapy based on tumour growth kinetics. Br J Cancer 105, 682–686 (2011).

25. Yamada, M., et al. Microscopic Venous Invasion in Pancreatic Cancer. Ann Surg Oncol 25, 1043–1051 (2018).

26. Oettle, H., et al. Adjuvant chemotherapy with gemcitabine vs observation in patients undergoing curative-intent resection of pancreatic cancer: a randomized controlled trial. JAMA 297, 267–277 (2007).

27. Sinn, M., et al. CONKO-005: Adjuvant Chemotherapy With Gemcitabine Plus Erlotinib Versus Gemcitabine Alone in Patients After R0 Resection of Pancreatic Cancer: A Multicenter Randomized Phase III Trial. J Clin Oncol 35, 3330–3337 (2017).

28. Rhim, A.D., et al. EMT and dissemination precede pancreatic tumor formation. Cell 148, 349–361 (2012).

29. Bulle, A., et al. Gemcitabine induces Epithelial-to-Mesenchymal Transition in patient-derived pancreatic ductal adenocarcinoma xenografts. Am J Transl Res 11, 765–779 (2019).

30. Ooft, S.N., et al. Prospective experimental treatment of colorectal cancer patients based on organoid drug responses. ESMO Open 6, 100103 (2021).

31. Herpers, B., et al. Functional patient-derived organoid screenings identify MCLA-158 as a therapeutic EGFR x LGR5 bispecific antibody with efficacy in epithelial tumors. Nat Cancer 3, 418–436 (2022).

32. Velez-Velez, L.M., Hughes, C.L. & Kasi, P.M. Clinical Value of Pharmacogenomic Testing in a Patient Receiving FOLFIRINOX for Pancreatic Adenocarcinoma. Front Pharmacol 9, 1309 (2018).

33. Chen, N., et al. Pharmacokinetics and pharmacodynamics of nab-paclitaxel in patients with solid tumors: disposition kinetics and pharmacology distinct from solvent-based paclitaxel. J Clin Pharmacol 54, 1097–1107 (2014).

34. Ciccolini, J., Serdjebi, C., Peters, G.J. & Giovannetti, E. Pharmacokinetics and pharmacogenetics of Gemcitabine as a mainstay in adult and pediatric oncology: an EORTC-PAMM perspective. Cancer Chemother Pharmacol 78, 1–12 (2016).

35. Rothenberg, M.L. Efficacy of oxaliplatin in the treatment of colorectal cancer. Oncology (Williston Park) 14, 9–14 (2000).

36. Catalano, M., Conca, R., Petrioli, R., Ramello, M. & Roviello, G. FOLFOX vs FOLFIRI as Second-line of Therapy After Progression to Gemcitabine/Nab-paclitaxel in Patients with Metastatic Pancreatic Cancer. Cancer Manag Res 12, 10271–10278 (2020).

37. Porter, R.L., et al. Epithelial to mesenchymal plasticity and differential response to therapies in pancreatic ductal adenocarcinoma. Proc Natl Acad Sci U S A (2019).

38. Evan, T., Wang, V.M. & Behrens, A. The roles of intratumour heterogeneity in the biology and treatment of pancreatic ductal adenocarcinoma. Oncogene 41, 4686–4695 (2022).

39. Matsuda, Y., et al. Objective assessment of tumor regression in post-neoadjuvant therapy resections for pancreatic ductal adenocarcinoma: comparison of multiple tumor regression grading systems. Sci Rep 10, 18278 (2020).

40. El Amrani, M., et al. Gemcitabine-induced epithelial-mesenchymal transition-like changes sustain chemoresistance of pancreatic cancer cells of mesenchymal-like phenotype. Mol Carcinog 58, 1985–1997 (2019).

41. Shen, Y., et al. miR-375 mediated acquired chemo-resistance in cervical cancer by facilitating EMT. PLoS One 9, e109299 (2014).

42. Aldonza, M.B.D., et al. Prior acquired resistance to paclitaxel relays diverse EGFR-targeted therapy persistence mechanisms. Sci Adv 6, eaav7416 (2020).

43. Werba, G., et al. Single-cell RNA sequencing reveals the effects of chemotherapy on human pancreatic adenocarcinoma and its tumor microenvironment. Nat Commun 14, 797 (2023).

44. Hahn, S., et al. Organoid-based epithelial to mesenchymal transition (OEMT) model: from an intestinal fibrosis perspective. Sci Rep 7, 2435 (2017).

45. Bates, R.C. & Mercurio, A.M. Tumor necrosis factor-alpha stimulates the epithelial-to-mesenchymal transition of human colonic organoids. Mol Biol Cell 14, 1790–1800 (2003).

46. Karve, K., Netherton, S., Deng, L., Bonni, A. & Bonni, S. Regulation of epithelial-mesenchymal transition and organoid morphogenesis by a novel TGFbeta-TCF7L2 isoform-specific signaling pathway. Cell Death Dis 11, 704 (2020).

47. Nasir, A., et al. A compromise between Tgfβ and Egfr signaling programs confers the ability to lead heterogeneous collective invasion. bioRxiv, 2020.2011.2014.383232 (2022).

48. Deben, C., et al. Expression of SARS-CoV-2-Related Surface Proteins in Non-Small-Cell Lung Cancer Patients and the Influence of Standard of Care Therapy. Cancers (Basel) 14(2022).

49. Le Compte, M., et al. Patient-derived organoids as individual patient models for chemoradiation response prediction in gastrointestinal malignancies. Crit Rev Oncol Hematol 157, 103190 (2021).

50. Hafner, M., Niepel, M., Chung, M. & Sorger, P.K. Growth rate inhibition metrics correct for confounders in measuring sensitivity to cancer drugs. Nat Methods 13, 521–527 (2016).

51. Gupta, A., Gautam, P., Wennerberg, K. & Aittokallio, T. A normalized drug response metric improves accuracy and consistency of anticancer drug sensitivity quantification in cell-based screening. Commun Biol 3, 42 (2020).

