## Supplemental figures for "Uncovering the hidden threat: single-organoid analysis reveals clinically relevant treatment-resistant and invasive subclones in pancreatic cancer"

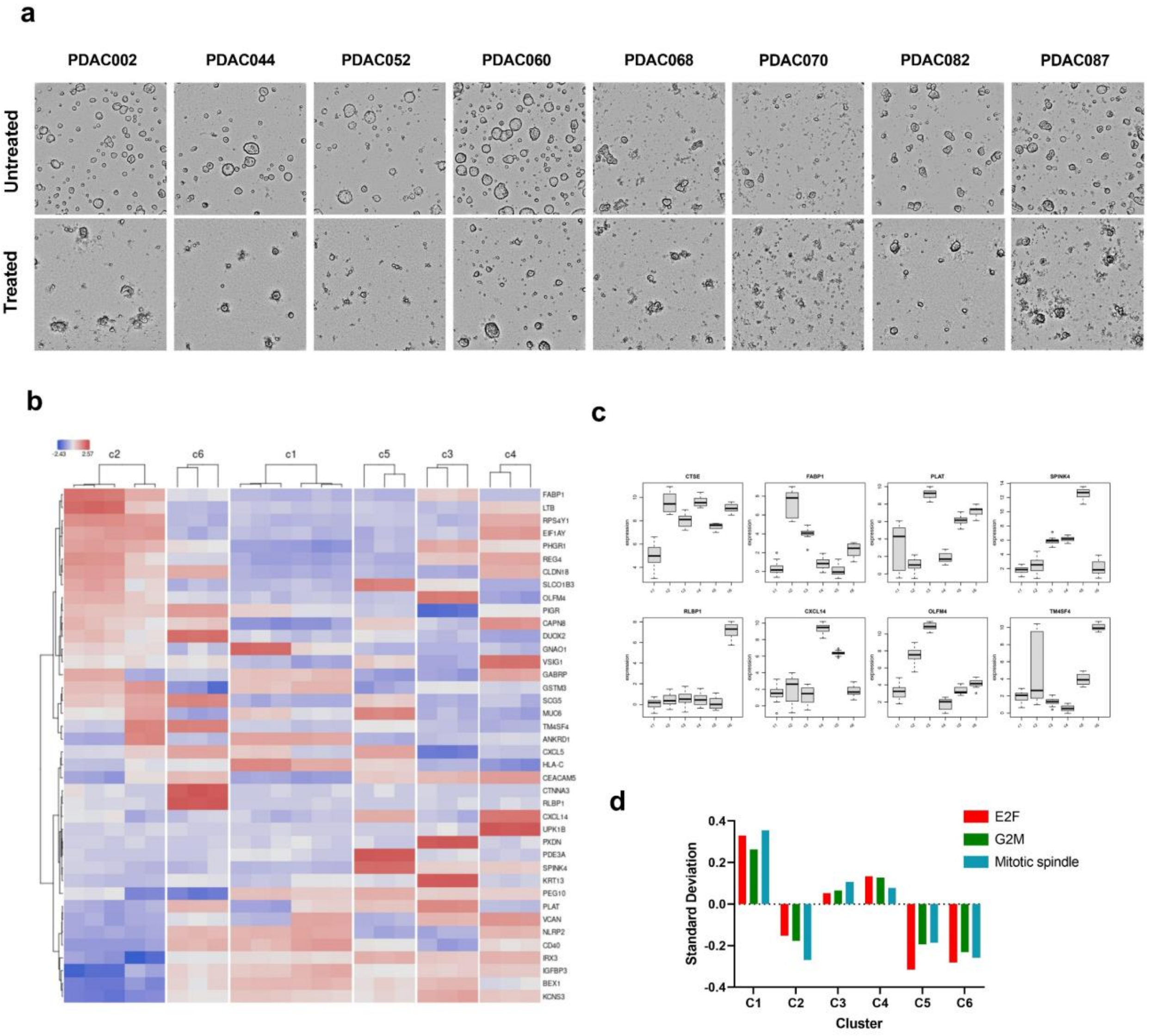
a
PDAC002
PDAC044
PDAC052
PDAC060
PDAC068
PDAC070
PDAC082
PDAC087
Untreated
Treated
b
-2.43
2.57
c2
c6
c1
c5
c3
c4
FABP1
LTB
RPS4Y1
EIF1AY
PHGR1
REG4
CLDN18
SLCO1B3
OLFM4
PIGR
CAPN8
DUOX2
GNAO1
VSIG1
GABRP
GSTM3
SCG5
MUC6
TM4SF4
ANKRD1
CXCL5
HLA-C
CEACAM5
CTNNA3
RLBP1
CXCL14
UPK1B
PXDN
PDE3A
SPINK4
KRT13
PEG10
PLAT
VCAN
NLRP2
CD40
IRX3
IGFBP3
BEX1
KCNS3
c
CTSE
FABP1
PLAT
SPINK4
RLBP1
CXCL14
OLFM4
TM4SF4
expression
d
E2F
G2M
Mitotic spindle
Standard Deviation
0.4
0.2
0.0
-0.2
-0.4
C1
C2
C3
C4
C5
C6
Cluster

**Extended Data Fig. 1 PDAC organoid cohort shows distinct morphological, clinical and molecular features. a.** Brightfield visualization of untreated and treated (gemcitabine-paclitaxel; 400 nM:80 nM) PDAC organoids showing the treatment-induces phenotypic variation. **b.** Overview of the top differently expressed genes between the individual clusters (C1,C2,C3,C4,C5 and C6). **c.** Bar-plot representation of potential cluster-specific biomarkers (most differently expressed genes between the clusters). d. Standard deviation (of the LogFC) of the selected Hallmark gene sets highlighting cluster specific growth/cell division features.

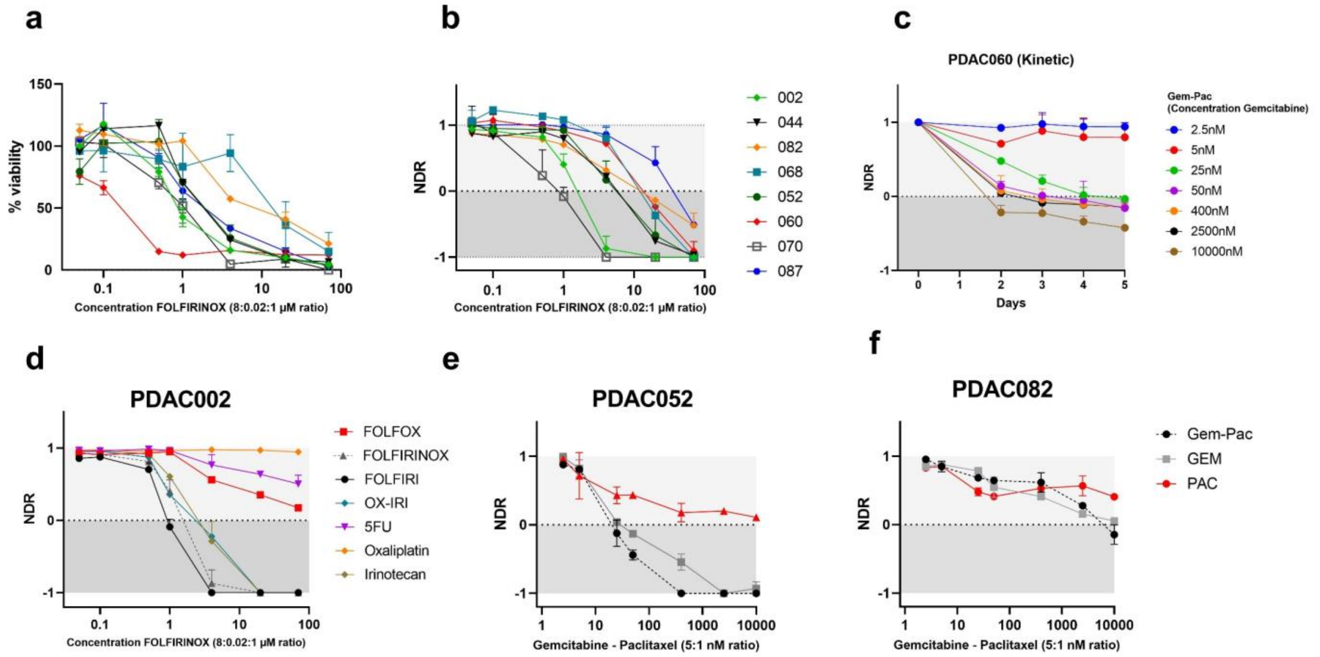
a
b
c
PDAC060 (Kinetic)
% viability
Concentration FOLFIRINOX (8:0.02:1 µM ratio)
NDR
002
044
082
068
052
060
070
087
Gem-Pac
(Concentration Gemcitabine)
2.5nM
5nM
25nM
50nM
400nM
2500nM
10000nM
Days
d
PDAC002
FOLFOX
FOLFIRINOX
FOLFIRI
OX-IRI
5FU
Oxaliplatin
Irinotecan
e
PDAC052
Gemcitabine - Paclitaxel (5:1 nM ratio)
f
PDAC082
Gem-Pac
GEM
PAC

**Extended Data Fig. 2 Normalized drug metric (NDR) highlights the inter-patient response heterogeneity. a-b.** % Viability and NDR dose response curve upon treatment with FOLRIRINOX (20 µM 5-FU:0.0625 µM SN38:2.5 µM Oxaliplatin ratio). **c**. Kinetic NDR quantification of PDAC060 treated with various concentrations (2.5 nM-10000 nM) gemcitabine-paclitaxel. NDR dose response curve of PDAC002 of the different FOLFIFINOX regimen. **e-f.** NDR dose response curve of PDAC052 (additive effect) and PDAC082 (no additive effect) showing the patient-specific additives effects.

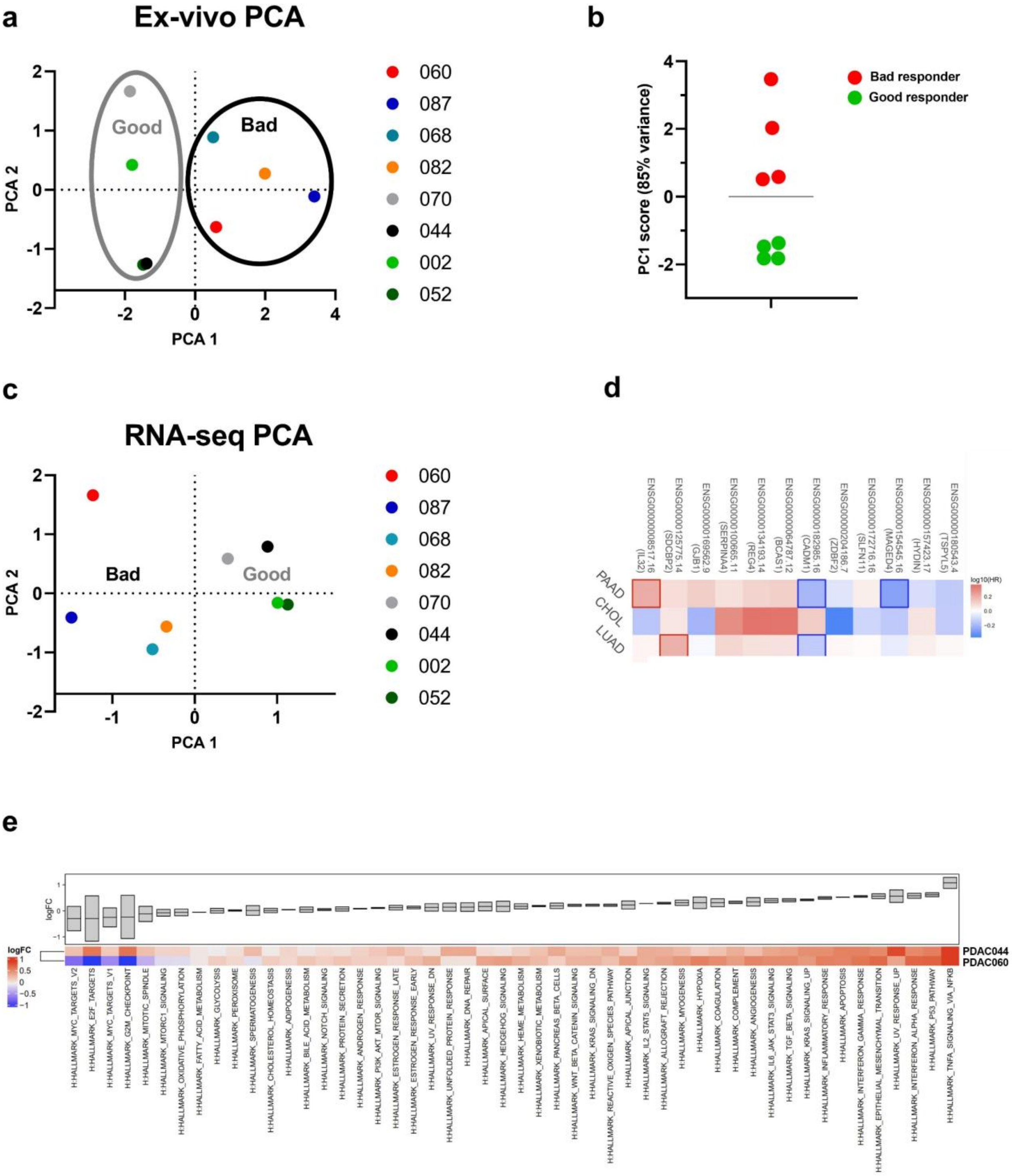
a
Ex-vivo PCA
Good
Bad
PCA 1
PCA 2
060
087
068
082
070
044
002
052
b
PC1 score (85% variance)
Bad responder
Good responder
c
RNA-seq PCA
Bad
Good
PCA 1
PCA 2
060
087
068
082
070
044
002
052
d
ENSG00000008517.16 (IL32)
ENSG00000125775.14 (SDCBP2)
ENSG00000169562.9 (GJB1)
ENSG00000100665.11 (SERPINA4)
ENSG00000134193.14 (REG4)
ENSG00000064787.12 (BCAS1)
ENSG00000182985.16 (CADM1)
ENSG00000204186.7 (ZDBF2)
ENSG00000172716.16 (SLFN11)
ENSG00000154545.16 (MAGED4)
ENSG00000157423.17 (HYDIN)
ENSG00000180543.4 (TSPYL5)
PAAD
CHOL
LUAD
log10(HR)
e
logFC
PDAC044
PDAC060
H:HALLMARK_MYC_TARGETS_V2
H:HALLMARK_E2F_TARGETS
H:HALLMARK_MYC_TARGETS_V1
H:HALLMARK_G2M_CHECKPOINT
H:HALLMARK_MITOTIC_SPINDLE
H:HALLMARK_MTORC1_SIGNALING
H:HALLMARK_OXIDATIVE_PHOSPHORYLATION
H:HALLMARK_FATTY_ACID_METABOLISM
H:HALLMARK_GLYCOLYSIS
H:HALLMARK_PEROXISOME
H:HALLMARK_SPERMATOGENESIS
H:HALLMARK_CHOLESTEROL_HOMEOSTASIS
H:HALLMARK_ADIPOGENESIS
H:HALLMARK_BILE_ACID_METABOLISM
H:HALLMARK_NOTCH_SIGNALING
H:HALLMARK_PROTEIN_SECRETION
H:HALLMARK_ANDROGEN_RESPONSE
H:HALLMARK_PI3K_AKT_MTOR_SIGNALING
H:HALLMARK_ESTROGEN_RESPONSE_LATE
H:HALLMARK_ESTROGEN_RESPONSE_EARLY
H:HALLMARK_UV_RESPONSE_DN
H:HALLMARK_UNFOLDED_PROTEIN_RESPONSE
H:HALLMARK_DNA_REPAIR
H:HALLMARK_APICAL_SURFACE
H:HALLMARK_HEDGEHOG_SIGNALING
H:HALLMARK_HEME_METABOLISM
H:HALLMARK_XENOBIOTIC_METABOLISM
H:HALLMARK_PANCREAS_BETA_CELLS
H:HALLMARK_WNT_BETA_CATENIN_SIGNALING
H:HALLMARK_KRAS_SIGNALING_DN
H:HALLMARK_REACTIVE_OXIGEN_SPECIES_PATHWAY
H:HALLMARK_APICAL_JUNCTION
H:HALLMARK_IL2_STAT5_SIGNALING
H:HALLMARK_ALLOGRAFT_REJECTION
H:HALLMARK_MYOGENESIS
H:HALLMARK_HYPOXIA
H:HALLMARK_COAGULATION
H:HALLMARK_COMPLEMENT
H:HALLMARK_ANGIOGENESIS
H:HALLMARK_IL6_JAK_STAT3_SIGNALING
H:HALLMARK_TGF_BETA_SIGNALING
H:HALLMARK_KRAS_SIGNALING_UP
H:HALLMARK_INFLAMMATORY_RESPONSE
H:HALLMARK_APOPTOSIS
H:HALLMARK_INTERFERON_GAMMA_RESPONSE
H:HALLMARK_EPITHELIAL_MESENCHYMAL_TRANSITION
H:HALLMARK_UV_RESPONSE_UP
H:HALLMARK_INTERFERON_ALPHA_RESPONSE
H:HALLMARK_P53_PATHWAY
H:HALLMARK_TNFA_SIGNALING_VIA_NFKB

**Extended Data Fig. 3 Transcriptional analysis highlights clinical relevance of NDR readout. a-c.** Principle components analysis (PCA) of either the ex-vivo based drug responses or transcriptome data (normalized read counts). **d**. Hazard ratio heatmap indicating that the 6-gene signatures can be extended towards other aggressive tumor types as well. PAAD= pancreatic ductal adenocarcinoma, CHOL=cholangiocarcinoma and LUAD=lung adenocarcinoma. **e.** Hallmark gene set enrichment analysis of PDAC044 and PDAC060 upon treatment with gemcitabine-paclitaxel (400 nM:80 nM).

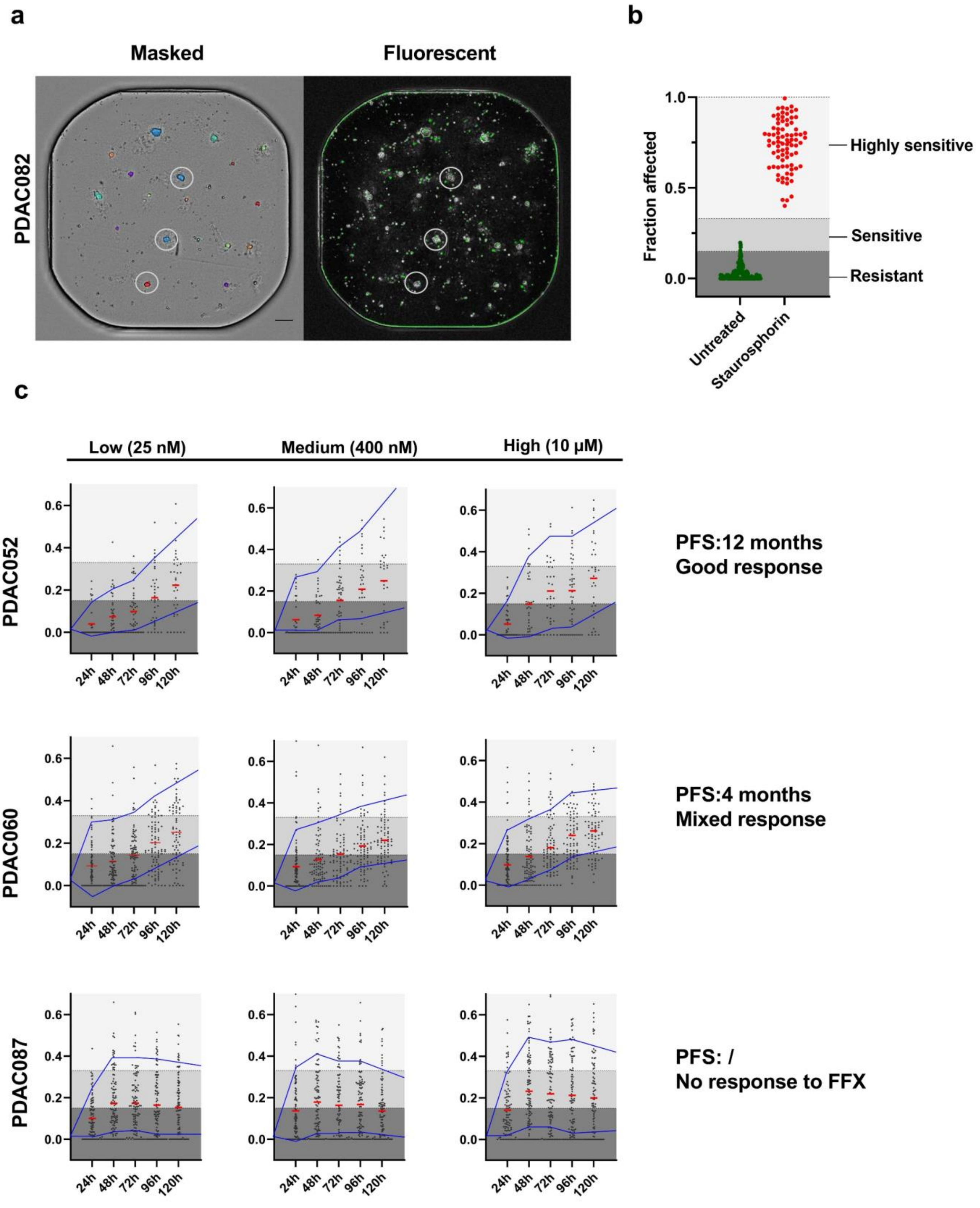
a
Masked
Fluorescent
PDAC082
b
Fraction affected
1.0
0.5
0.0
Highly sensitive
Sensitive
Resistant
Untreated
Staurosphorin
c
Low (25 nM)
Medium (400 nM)
High (10 μM)
PDAC052
PDAC060
PDAC087
24h
48h
72h
96h
120h
PFS:12 months
Good response
PFS:4 months
Mixed response
PFS: /
No response to FFX

**Extended Data Fig. 4 Single-organoid analysis reveals the intra-tumoral response heterogeneity. a.** Representative masked and fluorescent (cytotox green) images, showing the presence of masked PDAC organoid clones with a minor green overlap. For each masked organoids, the green overlapping area was automatically quantified (basis of the single organoid readout. Scale bar= 100 µM **b**. Fraction affected values of the control and positive control (staurosphorine), which was used to define the response ranges (sensitive, highly sensitive and resistant). **c.** Kinetic single-organoid quantification (fraction affected) upon treatment with low (25 nM:5 nM), middle (400 nM:80 nM) or high (10000 nM and 250 nM) concentrations of gemcitabine-paclitaxel.

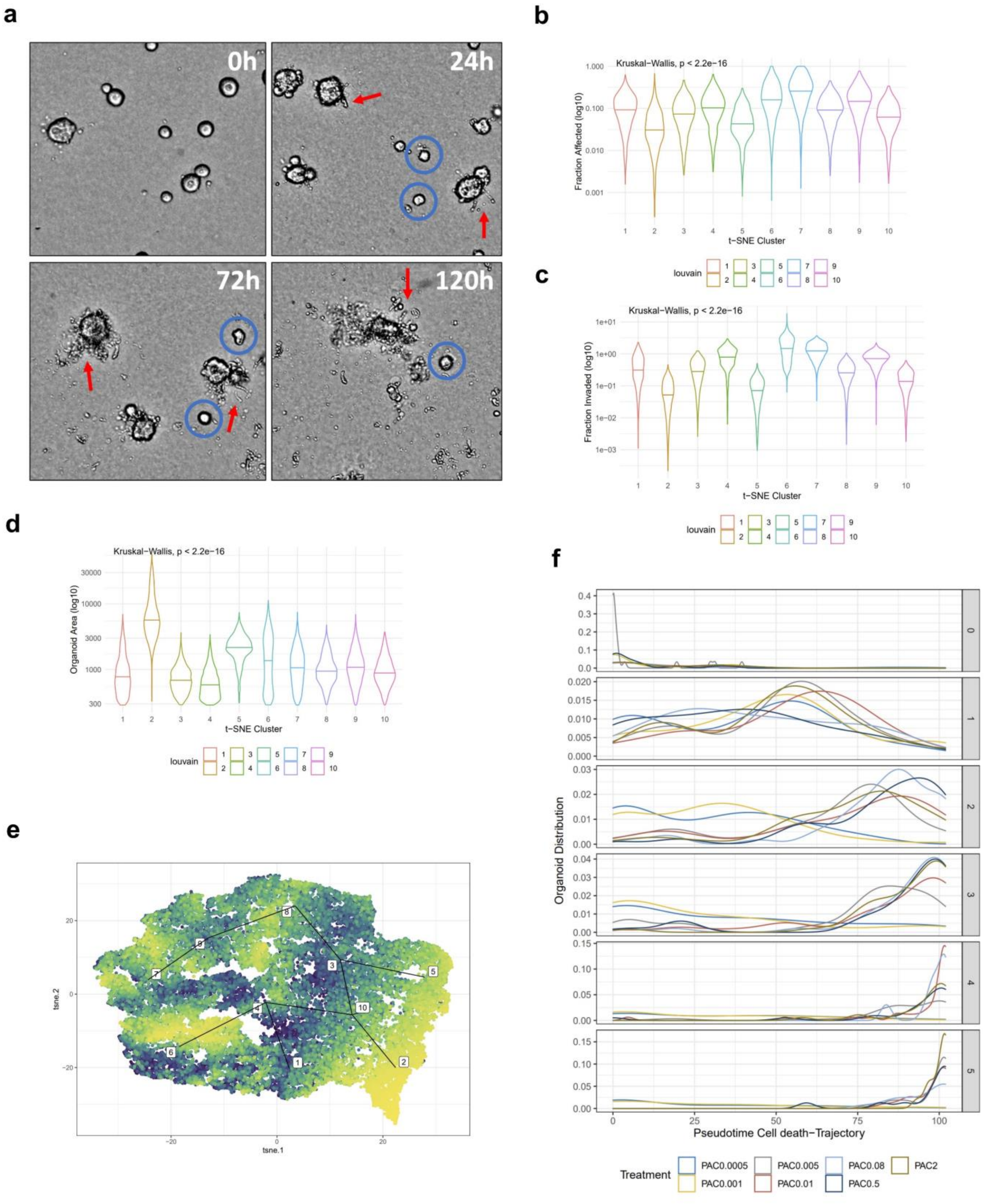
a
0h
24h
72h
120h
b
Kruskal−Wallis, p < 2.2e−16
Fraction Affected (log10)
t−SNE Cluster
louvain
c
Kruskal−Wallis, p < 2.2e−16
Fraction Invaded (log10)
t−SNE Cluster
louvain
d
Kruskal−Wallis, p < 2.2e−16
Organoid Area (log10)
t−SNE Cluster
louvain
e
tsne.2
tsne.1
f
Organoid Distribution
Pseudotime Cell death−Trajectory
Treatment
PAC0.0005
PAC0.001
PAC0.005
PAC0.01
PAC0.08
PAC0.5
PAC2

**Extended Data Fig. 5 PDAC organoids reveal patient-therapy-concentration and -time specific invasive patterns. a.** Brightfield visualization of PDAC002 showing the time dependent invasive behavior. Red arrows= spindle shaped cells, Blue circles=non-invasive PDAC organoids. Scale bar=20 µm. **b-d.** Specific features (Fraction affected, Invasive fraction and organoid area) per cluster. **e.** tSNE visualization of organoids with a high area (Cluster 2 and 5). **f.** Density distribution analysis over the Fraction affected (cell death) pseudotime trajectory.

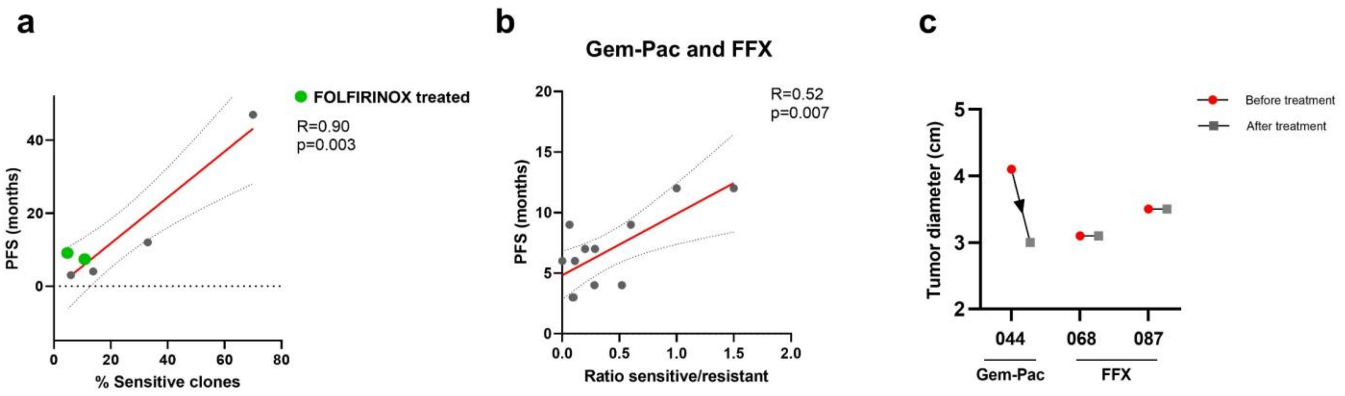
a
FOLFIRINOX treated
R=0.90
p=0.003
PFS (months)
% Sensitive clones
b
Gem-Pac and FFX
R=0.52
p=0.007
PFS (months)
Ratio sensitive/resistant
c
Before treatment
After treatment
Tumor diameter (cm)
044
068
087
Gem-Pac
FFX

**Extended Data Fig. 6 Clinical significance of NDR and single organoid analysis. a.** Correlation of the % sensitive, PDAC clones (4 µM 5-FU:0.0125 µM SN38:0.5 µM Oxaliplatin). **b.** Combined correlation of the ratio % sensitive/resistant clones of gemcitabine-paclitaxel and FOLFIRINOX (excluding PDAC044 to put the emphasis on the first 20 months). **c.** CT-based quantification of the tumor volume before and after treatment with Gemcitabine/Nab-Paclitaxel or FOLFIRINOX.
